# KIN: A method to infer relatedness from low-coverage ancient DNA

**DOI:** 10.1101/2022.10.21.513172

**Authors:** Divyaratan Popli, Stéphane Peyrégne, Benjamin M. Peter

## Abstract

Genetic kinship of ancient individuals can provide insights into their culture and social hierarchy, and is relevant for downstream genetic analyses. However, estimating relatedness from ancient DNA is difficult due to low-coverage, ascertainment bias, or contamination from various sources. Here, we present KIN, a method to estimate the relatedness of a pair of individuals from the identical-by-descent segments they share. KIN accurately classifies up to 3rd-degree relatives using ≥ 0.05*x* sequence coverage and differentiates siblings from parent-child. It incorporates additional models to adjust for contamination and detect inbreeding, which improves classification accuracy.

## 1 Introduction

### 1.1 Why study relatedness?

Identifying related individuals is a common task in genetic studies. Relatedness is of direct interest in e.g. DNA forensics, where familial search can aid in solving criminal cases, and to identify unknown deceased persons [29, 38]. Genetic paternity tests have an important application in resolving family relation, e.g. in establishing relationship between an individual applying for immigration and the claimed relatives [10]. It is also an essential step in population genetics and association studies, where samples are typically assumed to be independent random draws from the population. For animal and plant breeders and conservation biologists, reconstructing pedigrees and finding related individuals is important to avoid inbreeding and ensure diversity [14, 31, 16].

In ancient DNA studies, relatedness can be used to identify bones and teeth belonging to the same individual. Given adequate familiarity with the subject, relatedness can provide an understanding of an ancient society’s social structures, mobility and inheritance rules [1, 28, 40].

### 1.2 Approaches to estimate relatedness from high-coverage data

Commonly, pairs of related individuals are identified by looking for parts of the genome that are identical by descent (IBD), i.e. inherited from a recent common ancestor. Due to the laws of Mendelian segregation, each parent will share exactly one set of chromosomes IBD with their offspring, while subsequent recombination means that a grandparent will, on average share a quarter of their genome with a grand-child. Along the genomes of a pair of diploid individuals, there are three IBD states possible at any given position: the individuals share zero, one or two chromosomes IBD. The genome-wide proportions of these states (usually referred to as *k*_0_, *k*_1_, *k*_2_, so that *k*_0_ +*k*_1_ +*k*_2_ = 1) can be used to infer the degree and nature of relatedness for a pair of individuals. For example, a pair of siblings are expected to have all three possible IBD states with proportions of 0.25,0.5,0.25, respectively (Fig. 1). These IBD probabilities can directly be used to categorize their relatedness as shown in Table 1. One can also use these probabilities to estimate the coefficient of relatedness *r*, which is defined as the proportion of the genome that is IBD. In the absence of inbreeding, this would be calculated as *r* = *k*_1_/2 + *k*_2_.

**Figure 1:**
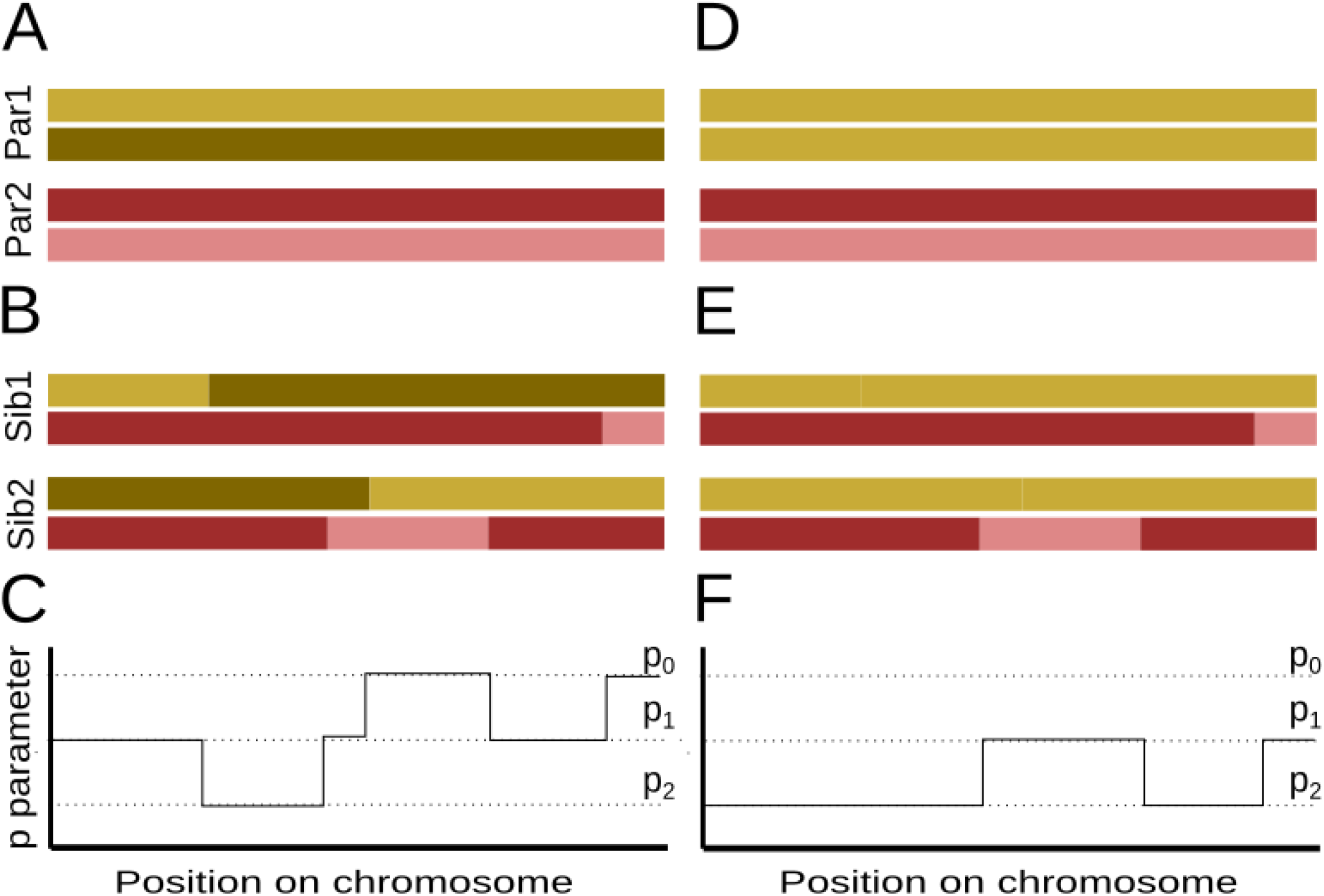
IBD sharing between siblings without and with runs of homozygosity (ROH). (A) Schematic of chromosomes for two parents (Par1, Par2). (B) Schematics of recombinant chromosomes of two children (Sib1, Sib2). (C) Expected differences between Sib1 and Sib2 (*p*) along the chromosome. In both cases, *p* can take values of *p*_0_, *p*_1_ or *p*_2_, which are the expected proportion of differences in IBD states 0,1 and 2, respectively. (D-F): Same as A-C, except parent 1 is assumed to be homozygous.

**Table 1:** IBD sharing probabilities for different relations in absence of inbreeding

| Relatedness | $k_0$ | $k_1$ | $k_2$ |
| --- | --- | --- | --- |
| Unrelated | 1 | 0 | 0 |
| 3rd Degree | 0.75 | 0.25 | 0 |
| 2nd Degree | 0.5 | 0.5 | 0 |
| Siblings | 0.25 | 0.5 | 0.25 |
| Parent-Child | 0 | 1 | 0 |
| Identical/twins | 0 | 0 | 1 |

However, since it is not possible to directly observe IBD segments, a common approach is to first identify segments of the genome that are Identical by State (IBS) and to use population allele frequencies obtained from an out-of-sample reference panel to calculate the probability of IBD given IBS. There are several methods that incorporate reference panel allele frequencies, phase information, recombination maps, or genotype calls to co-estimate IBD and the relatedness coefficient [15, 22, 43, 4, 25, 37, 27, 12, 30, 5, 23].

### 1.3 Approaches that address problems with ancient DNA data

One major issue with applying the above-mentioned methods to ancient DNA data is that the sequence coverage is typically low, making it difficult to obtain accurate genotype calls. Several methods surmount this problem by using genotype likelihoods [24, 20]. In this way, it is possible to account for the uncertainty in genotype calls by summing over all possible genotypes, weighted by their genotype likelihoods. However, these approaches typically still require at least 2x coverage, since genotype likelihoods may be imprecise at lower coverages [19] . Ancient DNA analyses often face additional challenges such as the unavailability of reference panels to estimate population allele frequencies, contamination with present-day DNA [32], and an ascertainment bias caused by DNA capture approaches [36, 44].

Several methods have been proposed to estimate relatedness without a reference panel from ancient DNA, but they require either > 4x coverage [46], or a large sample size to get an estimate of allele frequencies in the population from which the sampled individuals originate [42]. A second issue is contamination. If the contamination stems from another population, contaminated data will look more dissimilar to other individuals from the analyzed population, and hence relatedness will be underestimated. In addition, some analyzed genomes may have long runs of homozygosity (ROH), for example due to a small population size, or recent inbreeding. Long ROH cause related individuals to seem genetically more similar to each other, but do not affect the genetic distance between unrelated individuals.

Moreover, ancient DNA is commonly captured with a SNP array that enriches for informative variants. Particularly, methods based on the fraction of sites in different IBS states are sensitive to the ascertainment bias caused by this non-random selection of targeted sites [46].

READ [21] addresses several of the issues encountered in the analysis of ancient DNA. In particular, the lack of genotype calls is dealt with by randomly sampling alleles from each individual. A string of these alleles at each position (called pseudo-haploids) are then compared to other individuals to estimate average pairwise genetic distances, which in turn are used to infer relatedness. However, READ only infers the degree of relatedness, and only up to second degree.

### 1.4 How our method works

Here, we present KIN (Kinship Inference), a Hidden Markov Model (HMM)-based approach to estimate genetic kinship and IBD from low-coverage ancient DNA data. KIN can detect up to 3^rd^-degree relatives, and differentiates between siblings and parent-child relationships. KIN is also able to take into account ROH and contamination, and is not sensitive to SNP ascertainment. We validate the performance of KIN using simulations and show that we are able to infer relatedness in real data from two datasets: a group of Neandertals and a group of Bronze age individuals.

## 2 Algorithm

To infer relatedness, KIN fits one HMM for each pair of individuals and for each possible relatedness. The KIN-HMM infers IBD sharing between a pair of low-coverage individuals, optionally taking ROH tracts and contamination estimates in each individual into account. The best-fit model is then assigned as the inferred relatedness. If the locations of ROH tracts are unknown, we provide another HMM (ROH-HMM) to coarsely estimate the location of ROH for samples with sufficient coverage (≥ 0.1*x*). Our method is available on https://github.com/DivyaratanPopli/Kinship_Inference along with a python package (KINgaroo) to generate the input files for the models directly from bam files.

### 2.1 Model description

The goal of KIN-HMM is to infer how two individuals are related via the patterns of shared IBD states along a pair of genomes. For this purpose, we subdivide the genomes of a pair of individuals into *L* large genomic windows (typically of size 10Mb), and infer the pattern of IBD-sharing for each relatedness case we consider (Unrelated, 5^*th*^ Degree, 4^*th*^ Degree, 3^*rd*^ Degree, Grandparent-Grandchild, Avuncular, Half-siblings, Parent-Child, Siblings, and Identical). We then compare the likelihood between all models, and classify each pair to the model with the highest likelihood. We also return the most likely locations of IBD tracts (using standard Viterbi algorithm), and the IBD state posterior probabilities.

The details of the likelihood computation are given in section 2.3. As is the case for any HMM, KIN-HMM requires a set of emission (section 2.4) and transition probability matrices. We assume the transition matrix for each relatedness case is fixed. Our emissions include a vector of parameters (*δ*) that describe the variance in the data for each IBD state that we infer from the data (section 2.4).

### 2.2 Input of KIN-HMM

The inputs of our algorithm are i) the number of overlapping sites for the *w*-th window *N_w_* for which both samples have at least one read available, ii) the number of pairwise differences *D_w_* at these sites, and iii) the probability of ROH in windows, by default obtained from ROH-HMM described in section 2.7.

For high-coverage data, *D_w_* can be directly obtained by comparing genotypes, but for low-coverage samples, *D_w_* needs to be estimated from the sequencing data. The simplest approach is to randomly-sample a read from each position [13, 35, 11]. However, such an approach may result in loss of data, and hence we estimate *D_w_* by implicitly summing over all possible samplings:

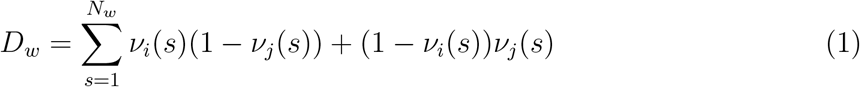

Here, *ν_i_*(*s*) and *ν_j_*(*s*) are the proportions of reads carrying the derived allele at SNP index *s* for individuals i and j respectively. Throughout, we will use bold-face notation to refer to the vector (or matrix) collecting all the terms, e.g. **D** = (*D*_1_, *D*_2_, . . . *D_L_*).

### 2.3 Log likelihood of the KIN-HMM

The KIN-HMM uses **D** and **N** to classify each window into three hidden states *Z_w_* ∈ (0, 1, 2), reflecting zero, one or two shared chromosomes IBD, respectively. To take ROH into account, we define the variable *H_w_* ∈ (0, 1, 2) that designates that zero, one or both individuals are homozygous in window *w*. Since *H_w_* is unobserved, in practice we use the estimates from ROH-HMM: *h_wj_* = *P* (*H_w_* = *j*).

There are three additional model parameters, ***π***, **A** and ***δ***. The transition matrix **A** gives the probability of moving from state *i* to state *j*, given by *a_ij_*, and is fixed and estimated from simulations for each relatedness case (section 5.1). The initial probabilities ***π*** give the probabilities of being in each state *Z*_0_ (at the beginning of each chromosome), which we set to the stationary distribution of the transition matrix for simplicity. The overdispersion-parameter ***δ*** takes into account that SNPs in each window vary in their allele frequency (see next setion). For compactness of notation, we group the fixed parameters: *θ* = (**N**, **A**, ***π***).

Thus, the complete data likelihood for the HMM is

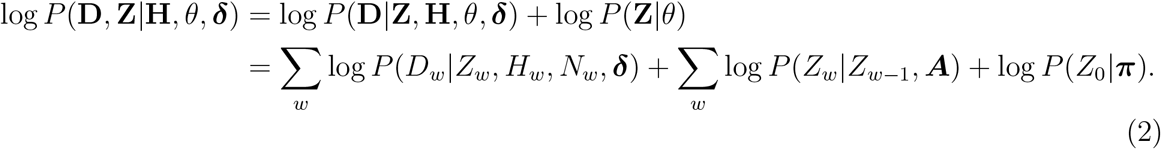

Here, **Z** is not dependent on **H** and ***δ***.

### 2.4 Emission probability

Using this setup, we can isolate the emissions *P* (*D_w_*|*Z_w_*, *H_w_*, *θ*, ***δ***) from equation 2 and optimize them for ***δ***. The simplest model is to assume that sites in each window are equally distributed and independent, in this case we could use the binomial likelihood:

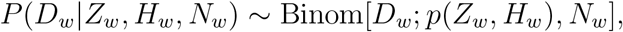

where, *p* is the proportion of differences expected for a particular IBD and ROH state. If the two individuals are unrelated in a particular window (i.e. *Z_w_* = 0), then the expected proportion of pairwise differences depends solely on the population history, and we denote this proportion with *p*_0_. If the two individuals share one or even both copies of the genome IBD, we would expect the proportion of differences to be reduced to 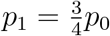, and 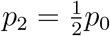, respectively, since either one or two of the four possible comparisons will be between identical chromosomes [21]. Thus, *p*(*Z_w_* = *i*, *H_w_* = 0) = *p_i_*.

The proportion of differences between unrelated individuals *p*_0_ is an important parameter. We follow READ [21] and estimate *p*_0_ as the median of differences for all possible pairs of individuals, which works well if the majority of individuals in the sample are unrelated.

The presence of long tracts of homozygosity resulting from recent inbreeding adds an additional complication, as the number of shared chromosomes may be overestimated [47]. For example, when considering two bones from the same individual, we would expect the entire genome to have a pairwise difference of *p*_2_, because two out of the four compared chromosomes are identical copies. However, in inbred regions all four chromosomes will be identical, and so the expected pairwise differences are zero (*p*_4_ in equation 3). Note however, that *p*_0_ does not depend on the presence of ROH, since all comparisons are between unrelated chromosomes even if both individuals are homozygous at a particular locus.

Taken together, we can summarize ***p*** in the following matrix, where rows give the state of *Z_w_*, and columns of *H_w_*:

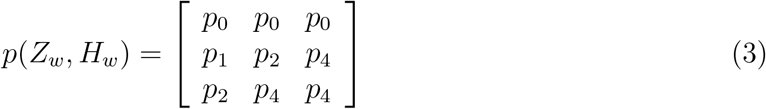

As explained above, we would expect *p*_4_ to be zero. However, as we do all our calculations in large windows, the start/end positions of windows may not coincide with that of ROH tracts, and we found that we obtain better results by setting 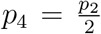, to take into account that many windows will only partially have four comparisons between identical chromosomes.

The effect of these considerations is that even though we have nine possible combinations of *Z_w_* and *H_w_* for each window, there are actually only four unique *p*-parameters *p_i_* with *i* ∈ (0,1,2,4).

#### Beta Binomial Model

We empirically find that the data often has considerably higher variance than would be expected from a binomial model (Fig. S2). We take this into account by adding an over-dispersion parameter ***δ***. Just like *p*(*Z_w_, H_w_*), *δ*(*Z_w_, H_w_*) depends on the number of chromosomes compared, and so each of the four *p_i_* has a corresponding *δ_i_* parameter.

Taken together, our emission probabilities are

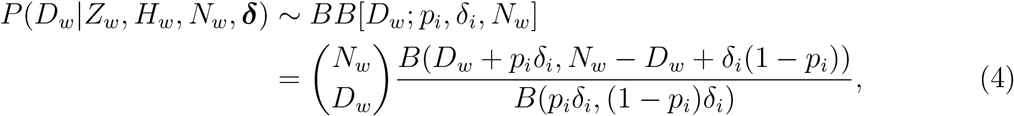

where *i* is fully determined by the combination of *Z_w_* and *H_w_* (see equation 3).

This parameterization of the beta distribution in terms of expected value *p* and over-dispersion *δ* is also called the Balding-Nichols-model [2], and is distinct from the more common parameterization in terms of *α* and *β*. We use this equation even if preprocessing steps (see equation 1 and section 2.8) result in non-integer *D_w_* and *N_w_*, in which case we approximate the binomial coefficient using Gamma functions.

### 2.5 Estimation of *δ*

We estimate the *δ*-parameters using an Expectation-Maximization (EM) algorithm [7].

#### Initialization

The value of *δ_i_* is unknown to start with, and we set it to a random value between 0 and 1000.

#### Expectation step

In the *t*-th iteration, we calculate the posterior probability of each IBD state in each window 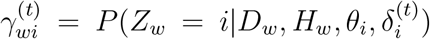 using the forward-backward algorithm, where 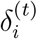 is the current estimate for *δ* for a given IBD state.

#### Maximization step

The only free parameters we estimate in the maximization step are the over-dispersion parameters *δ_i_*. We do this optimization using a cost function, which is the log-emission probability weighted by the posterior probabilities of the hidden states *γ_wj_* and optionally the ROH state-probabilities *h_wω_* obtained from the ROH-HMM.

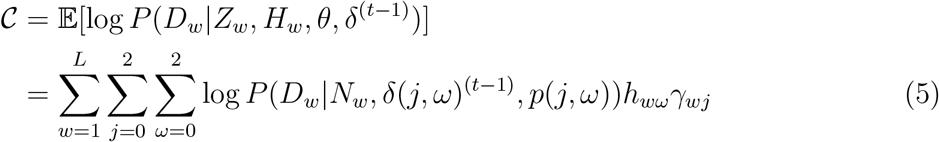

Using equation 3, we simplify this by grouping all the terms that have the same number of pairwise comparisons between identical chromosomes, which would result in the same *p_i_* and *δ_i_*, i.e.

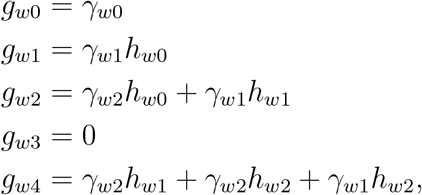

where *g*_*w*3_ is always 0 because there is no case that leads to only three comparisons between identical chromosomes.

So we can rewrite equation 5 as:

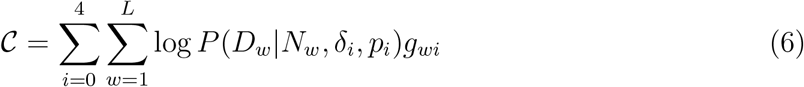

The cost function (equation 6) has one independent term for each *i* and so we can separate them and estimate each *δ_i_* independently using the minimize scalar algorithm implemented in `scipy.optimize` [45].

We constrain the optimization space of the *δ_i_*, as unconstrained optimization could result in some confounding of cases. We know that different cases of relatedness have different numbers of IBD states possible. For example, siblings may have all three IBD states present while a Parent-Child has only *Z_w_* = 1. However, the parent-child model could fit data generated under the sibling model by assigning it a very high *δ*_1_, which would reduce performance (for example, see Fig. S5, S6). We avoid this problem by constraining the ***δ*** such that the beta distributions for the different relatedness cases overlap by at most one standard deviation.

### 2.6 Model comparison

To infer the most likely relatedness case, we run our model on all relatedness cases mentioned in section 2.1, and compare the resulting likelihoods. We output the relatedness corresponding to the maximum likelihood model. Since we compare models where the parameters are not subset of each other, standard likelihood-ratio theory for nested models cannot be used to obtain confidence intervals. Instead we use the log likelihood ratio between the two best models as a statistic to assess the confidence in our classifications, and use simulations to obtain critical values (Fig. S8).

#### Grouping of cases

Particularly for low-quality data, we may not be able to distinguish all cases. Thus, we group 4^*th*^ and 5^*th*^ degree relatives together with unrelated. Similarly, we group half-siblings, avuncular and grandparent-grandchild to 2^*nd*^-degree relatives in the final results. We report the final pairwise classification in the following categories: Unrelated, Third Degree, Second Degree, Parent-Child, Siblings, Identical individuals.

#### Critical Values

To investigate the limits of our method, we plotted the true-positive and false-positive rates for classification of different relatedness in control simulations (without contamination, ROH and ascertainment bias, see section 5.1) when we use a particular difference in log-likelihood as a cutoff (Fig. S8). The figure shows that for all relatedness cases except for 3^*rd*^-degree, the false positive rate is below 5% even when simply selecting the model with the highest likelihood. We observe that for 3^*rd*^-degree relatives, using a cutoff of 1.0 brings down the false positive rate close to 5% for all coverages except 0.05x. Thus, we recommended using a cutoff of 1 for all cases where ROH and contamination are not a concern.

#### Example case of siblings

In Fig.2 we show the inferred IBD fragments from different KIN-HMMs when they are applied to simulated data from a pair of siblings. The models for identical, parent-child or unrelated relationships allow for just one IBD state, resulting in a flat line and low likelihood for this data. The other three cases all allow for different IBD-states, but the siblings-model predictions match true IBD states the most, as reflected by the highest log-likelihood and the close correspondence of the inferred and true IBD states.

**Figure 2:**
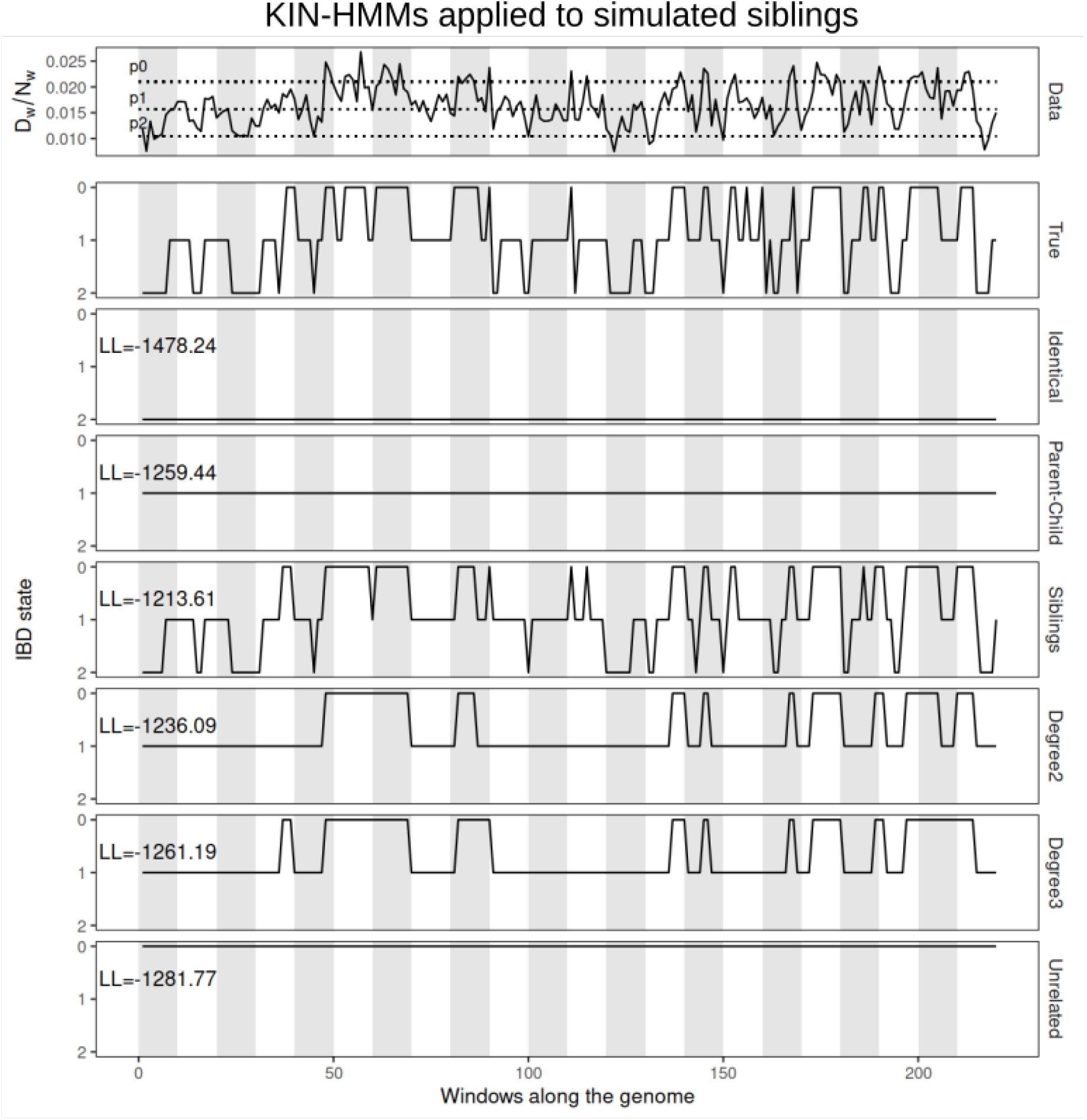
Comparison of pairwise difference data and inferred IBD fragments. The top panel shows the proportion of differences in each window along the genome for a pair of simulated siblings. Dashed lines represent *p*_0_, *p*_1_ and *p*_2_ estimates. The second panel shows the true IBD state for each window. The remaining panels show the IBD states predicted by particular relatedness models. The log-likelihood value for each model is shown on upper left corner of the panel. Light and shaded backgrounds represent distinct chromosomes.

### 2.7 ROH estimation model

Our HMM to detect ROH tracts works similarly to the KIN-HMM described above, but in this case we only consider one individual at a time, and only consider positions covered by at least two reads. For each site, we calculate the proportion of reads that carry different alleles, and sum them up in windows along the genome. We call the vector with the number of differences **∆**, and the vector with the number of sites with at least two reads **M**. Our model has two possible hidden states: homozygous state (*Y_w_* = 4), and non-homozygous state (*Y_w_* = 2). As above, we collect the hidden states in a vector **Y** = (*Y*_1_, . . . *Y_w_*, . . ., *Y_L_*). The complete data likelihood for the model in this case is then:

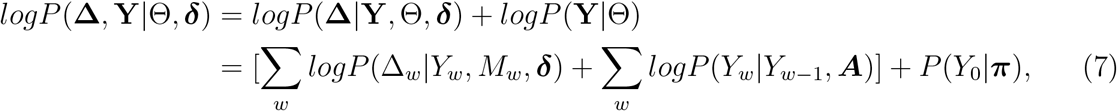

where Θ is a vector of initial probabilities (***π***), transition matrix (***A***) and ***M***.

Since the source of ROH may not be known, we estimate both transitions and emissions. We calculate the emissions using a beta-binomial likelihood, and fix the mean of the distributions corresponding to *Y_w_* = 4 and *Y_w_* = 2 at expected proportion of differences in a homozygous tract (*p*_4_) and expected proportion of differences in a non-homozygous tract (*p*_2_) respectively. The expectation step outputs the posterior probability Γ of being in state *Y_w_* = 4 or the state *Y_w_* = 2 in each window. The maximization step for emissions is analogous to that in the KIN model (eq. 5), and the optimization step here is done with the following cost function:

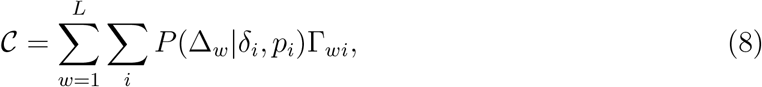

where *P*(∆_*w*_|*δ_i_*, *p_i_*) is a beta-binomial probability with mean *p* and over-dispersion parameter *δ* similar to eq. 4, and *i* can take values 2 and 4 corresponding to the hidden states *Y_w_*.

To estimate transitions, we initialize the transition matrix with the value 0.2 for the off-diagonal entries, and update it using the standard Baum-Welch update step [3].

Similar to the KIN-HMM, we avoid fitting issues by forcing all windows whose proportion of differences is larger than *p*_2_ to be in the non-homozygous state (Fig. S7).

### 2.8 Contamination correction

Contamination by DNA from present-day people is a common feature of human ancient DNA datasets [33]. To address this issue, we developed a heuristic that adjusts both *D_w_* and *N_w_* to minimize the influence of contamination on the relatedness inference.

We assume that contamination rates in both individuals are known and small (< 5%), and set *C_ij_* = *C_i_* +*C_j_*, where *C_i_,C_j_* are the contamination estimates from the two individuals. We also assume the divergence *ϕ* between our target population and the putative contaminant popoulation is known. With probability *C_ij_*, a comparison between two random reads from the pair of tested individual will contain a contaminant read, and thus contain a difference with probability *ϕ*, and with probability 1 − *C_ij_* it will be between endogenous ones. The comparisons between two contaminant reads are ignored, since we assume *C_ij_* to be small.

We estimate the expected number of differences from comparison of endogenous reads 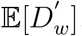, and the total number of sites with overlapping endogenous reads 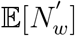.

For any particular comparison showing a difference *D* (not to be confused with the number of differences *D_w_*), we calculate the probability of the event *E* that it is between endogenous reads as

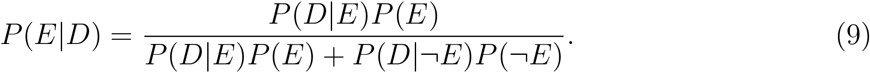

Then, by linearity of expectation, we obtain our estimator for the expected number of endogenous comparisons with a difference as

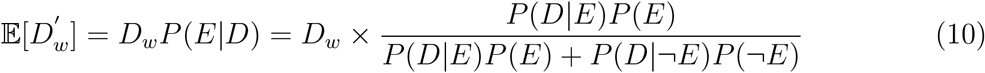

Of these terms, *P*(*E*) = 1 − *C_ij_* = 1 − *P* (¬*E*), and *P*(*D*|¬*E*) = *ϕ*.

For *P*(*D*|*E*), we use an estimator based on the genome-wide average:

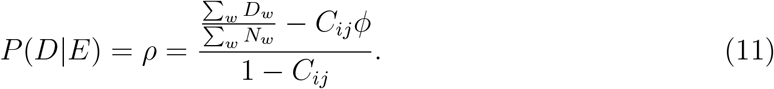

Taken together,

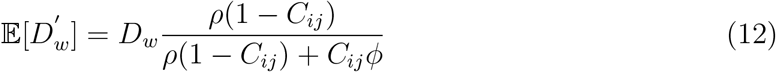

Analogous considerations lead to the expected number of endogenous comparisons that yield no difference:

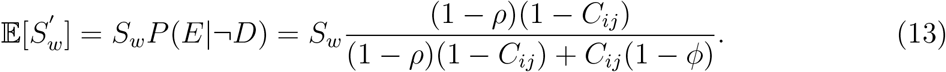

Here, *S_w_* = *N_w_* − *D_w_*. Hence, we set 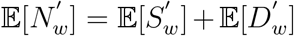. We do a similar contamination correction for the input of ROH-HMM.

## 3 Results

### 3.1 Evaluation with simulations

We first tested the performance of KIN with simulated pedigrees. We performed coalescent simulations to generate 8 unrelated diploid genomes and artificially mated them to form pedigrees of 17 individuals with relationships up to 5^*th*^ degree (Fig. S3). To evaluate the effect of sequence coverage on the performance of KIN, we generated artificial pileups at each polymorphic site for each individual following a Poisson distribution with 6 different average depths varying between 4*x* and 0.03*x* (see section 5). To mimic ROH, for some pedigrees, we picked a single allele at heterozygous sites in some regions as determined by a Markov chain so that on an average about 17% of the genome is ROH. We also created versions with contamination, by introducing alleles from distantly related individuals, and ascertainment bias, by selecting polymorphic sites identified in a subset of the individuals (see section 5). We created 60 pedigrees for each combination of average coverage and scenarios of presence/absence of ROH, contamination, and ascertainment bias, totalling 2,880 pedigrees.

#### ROH detection

In Fig. 3, we present an example of data and inference of the ROH-HMM for simulations with and without ROH, potentially with ascertainment bias and contamination. In all cases, we find that the inferred ROH closely matches the simulations, but the confidence in the classification tends to increase with the simulated coverage.

**Figure 3:**
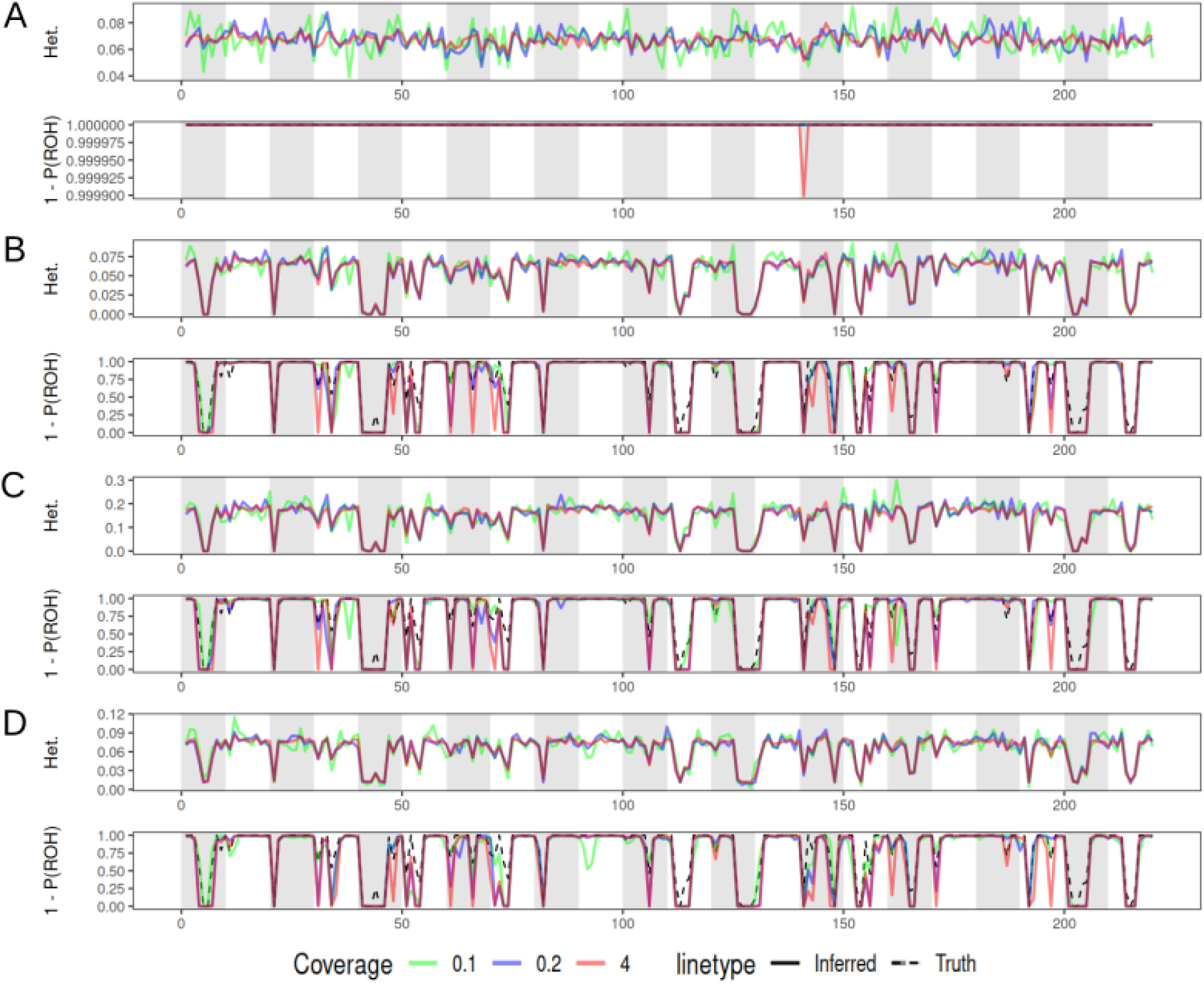
Estimation of ROH probabilities along the genome in simulations. The top row in each panel shows the proportion of differences in a simulated individual along the genome. In the bottom row, the dotted line shows the proportion of each window in ROH, and the solid lines show the estimated probability of not being in the ROH state. (A) Simulation with no ascertainment, contamination, or ROH. (B) Simulation with ROH. (C) Simulation with ROH and ascertainment. (D) Simulation with ROH and contamination.

A systematic evaluation of the performance of the ROH-HMM is given in Table 2: For the purpose of this analysis, we classified all windows that were at least 20% homozygous as ROH, and the remainder as non-ROH. Likewise, we classified all windows with a posterior probability of ROH of at least 20% as ROH. We used these cases to compute sensitivity (Se) and specificity (Sp). In the control case (simulations with no ascertainment, contamination, or ROH) we see that the specificity remains ≥ 0.99 for all coverages. In the cases where we simulate ROH, we find that sensitivity decreases from 97% at 4x coverage to less than 65% at 0.05x coverage, but specificity remains high at above 0.96 for all cases, suggesting that the number of erroneously called ROH segments is low.

**Table 2:**
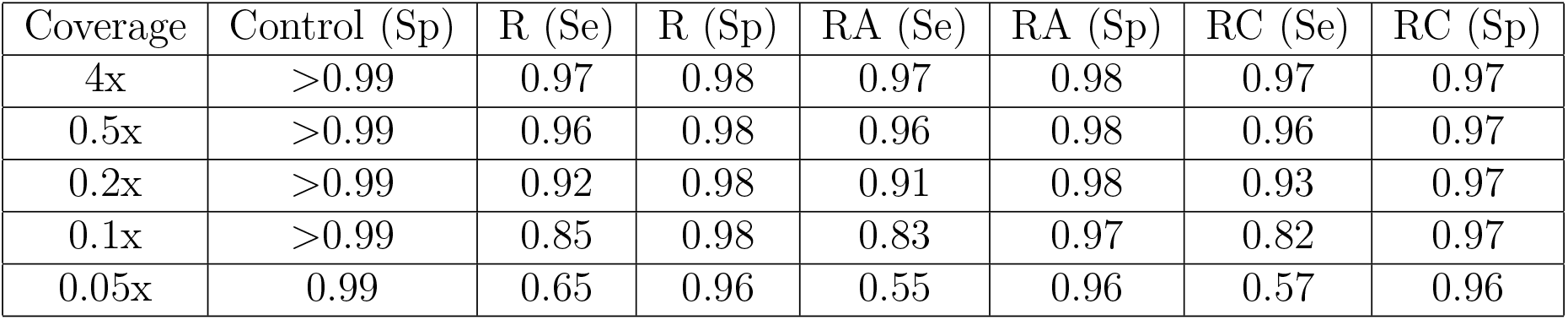
Model performance for ROH prediction. Here we test ROH-HMM in four different cases of simulations: Control (without ROH, ascertainment bias or contamination), R (with ROH), RA (with ROH and ascertainment bias), RC (with ROH and contamination)

#### IBD prediction

We investigated the accuracy of IBD state prediction along the genome by counting the number of genomic windows where we correctly predict IBD state for different relatedness and coverages (Fig. 4). The accuracy for coverages of 0.1x or higher is consistent and varies between 1.0 and 0.78, depending on relatedness. However, the accuracy decreases at lower coverages for most relatedness cases and ranges from 1.0 to 0.48 at 0.03x. The exceptions are for identical individuals and parent-child pairs, where the accuracy is nearly perfect for all investigated coverages, even at 0.03x coverage. We find that adding contamination, ascertainment bias in our simulations has little effect on IBD prediction. Adding ROH to the simulations reduces the IBD prediction in two cases: the average accuracy for second degree decreases from 0.89 to 0.85 and for siblings from 0.81 to 0.70 (Fig. S9). We see this adverse effect of ROH in case of siblings, perhaps because only in case of siblings we do not introduce long ROH directly, but long and short ROH may be present through parents (see section 5). Unidentified short ROH may cause a difficulty in differentiating between different combinations of IBD (*Z_w_*) and ROH states (*H_w_*). However, this does not affect the power to identify siblings in presence of ROH even at 0.05*x* (see Fig. 5)

**Figure 4:**
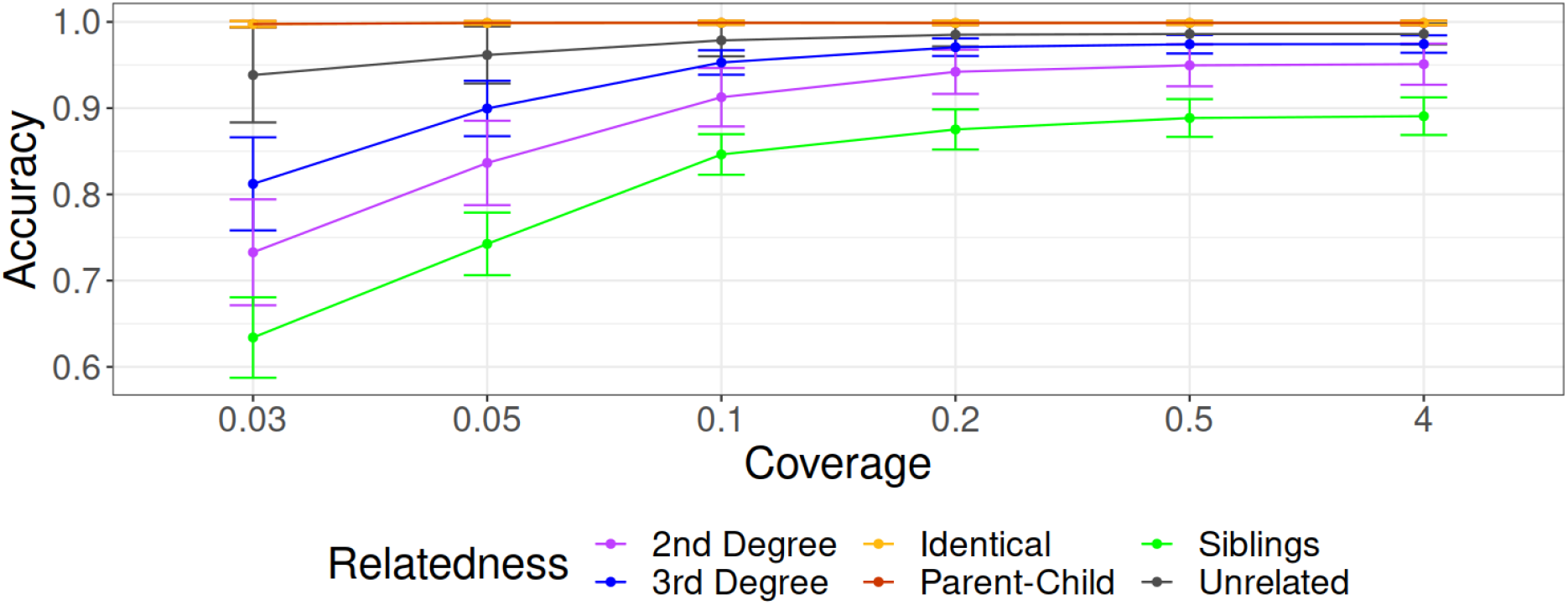
Evaluation of IBD estimation at different coverages. y-axis shows accuracy calculated over 60 simulations with each relatedness case and six different coverages shown on x-axis. The error bars are drawn at 1 standard deviation from the mean. Here, accuracy corresponding to relatedness cases for Parent-Child and Identical individuals is always 1, and overlaps with each other. Accuracy is defined as the proportion of correctly predicted IBD states, when compared to the central position of the window.

**Figure 5:**
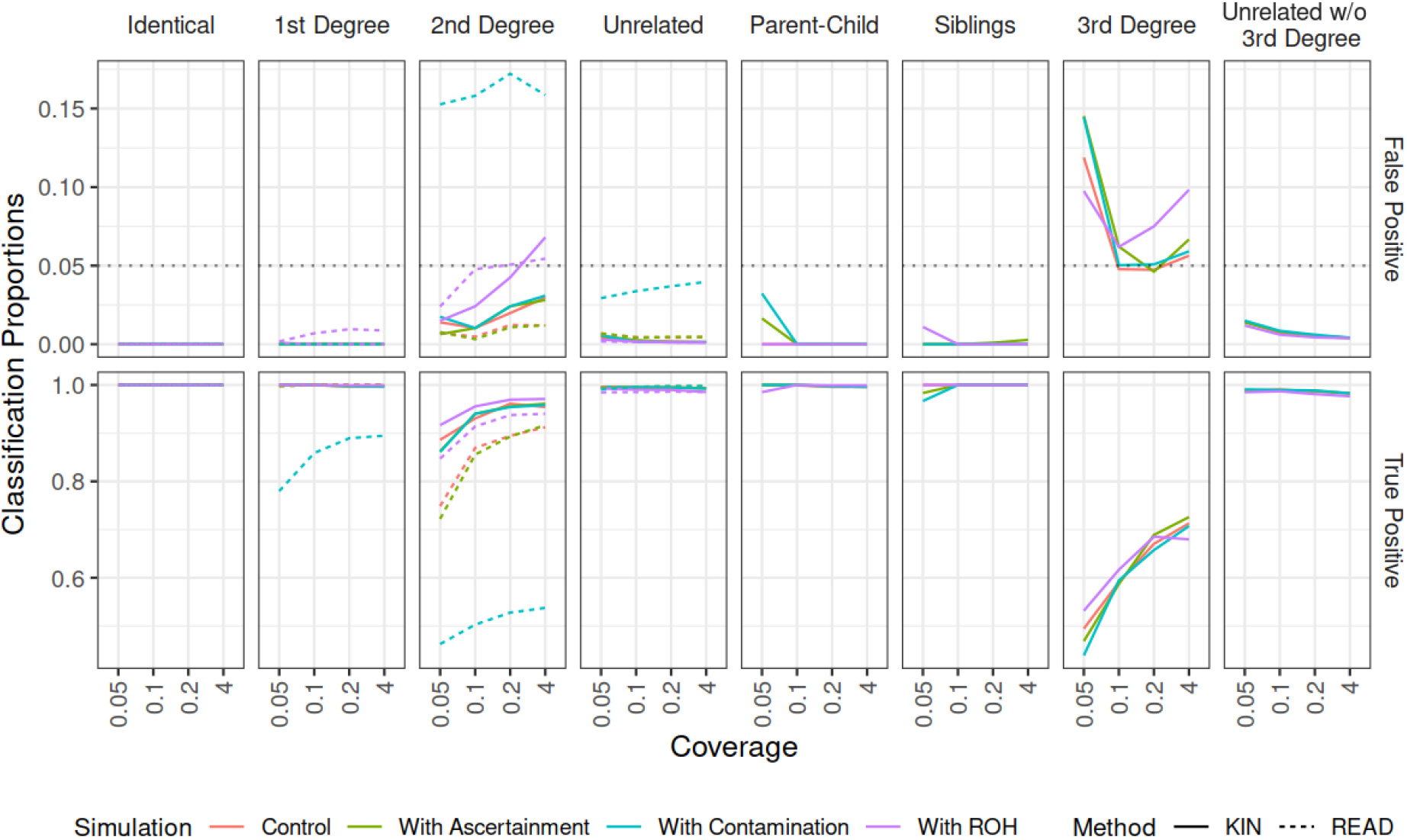
Comparison of KIN with READ using simulations with different coverages, and different cases of ascertainment, contamination and ROH. Unrelated label here refers to KIN performance results when all Unrelated, Fifth Degree, Fourth Degree, Third Degree pairs are labelled as Unrelated (for fair comparison with READ). ’Unrelated w/o 3^*rd*^ Degree’ refers to the performance results when 3^*rd*^ Degree is classified separately from the unrelated individuals.

#### Relatedness classification

We evaluated the classification accuracy of KIN (cutoff: 1 log-likelihood unit) and compared it to that of READ (cutoff: 1 standard deviation)) (Fig. 5). We first describe the results for relatedness cases detectable by READ, viz. identical, 1^*st*^ degree, 2^*nd*^ degree and unrelated. In this case we show that both methods have similar performance for low-coverage shotgun data (“control”-case), and for ascertained data. The true positive rate is above 0.97 for both KIN and READ, while the false negative rate is below 0.02. One exception is 2^*nd*^-degree relatives, where KIN has higher power than READ, and as coverage decreases from 4x to 0.05x, KIN’s true positive rate decreases from 0.95 to 0.89, compared to range of 0.91 to 0.75 for READ. The false positive rate in this case remains below 0.03 for both methods.

To investigate the impact of contamination, we performed simulations where we added up to 3% contamination to some individuals. We also find that READ is strongly impacted by contamination as the true positive rate is in the range 0.89 to 0.78 and 0.54 to 0.46 for 1st and 2nd degree relatives, respectively, and the false positive rate reaches up to 0.04 and 0.17 for unrelated individuals and 2nd degree relatives, respectively. In comparison, we also ran KIN, giving it the simulated contamination amounts for each individual. We found that the correction implemented in KIN is sufficient to remove this bias, and the true and false positive rates remain as in the control (Fig. 5). We also perform an analysis where we misspecify the contamination, and find that the effect of modest misspecification is small (Fig. S13).

For simulations with ROH, we find that KIN also outperforms READ for 1st and 2nd degree relatives, although both the true and false positive rates increased for both methods compared to the control for 2nd degree relatives. The increase in false positives is likely due to ROH making related individuals more similar. Finally, we also found that KIN has good power to detect relatedness cases that are not detectable by READ, i.e. parent-child, siblings and, to a lower level, 3rd degree relatives (Fig. 5).

### 3.2 Application to real data

#### Chagyrskaya and Okladnikov Neandertals

To test KIN on real ancient data, we applied it to a Neandertal dataset from Chagyrskaya and Okladnikov Caves in Siberia, Russia [18, 26, 41]. This dataset contains genetic data from a total of 16 skeletal remains that likely belong to contemporary Neandertals who occupied the Chagyrskaya and Okladnikov caves between 59 and 51 kya, and at least 44 kya respectively [41]. DNA extracted from each of these remains were captured with an array targeting variable sites identified in high coverage Neandertal and Denisovan genomes, and common variations in Africans [41]. This genetic data has low-to-intermediate depth of coverage ranging from 0.01x to 12.34x, with 8 samples at < 1*x* coverage. Some of these specimens showed signs of long ROH, and DNA contamination from modern humans as well as hyenas [41]. We focused our analysis on the variable sites in two high-coverage Neandertal genomes: Altai Neandertal (Denisova 5) [35] and Vindija 33.19 [34] as done by the authors for the relatedness analysis [41]. Our results for pairwise relatedness for these individuals are shown in Fig.6. We found three specimens from the same individual (Chagyrskaya13-Chagyrskaya19-Chagyrskaya1141), a parent-child pair (Chagyrskaya07 and Chagyrskaya17), and a pair of 2^*nd*^-degree relatives (Chagyrskaya01/Chagyrskaya60). Further, we identify Chagyrskaya17 and Chagyrskaya60 as 3rd-degree relatives (table S1).

**Figure 6:**
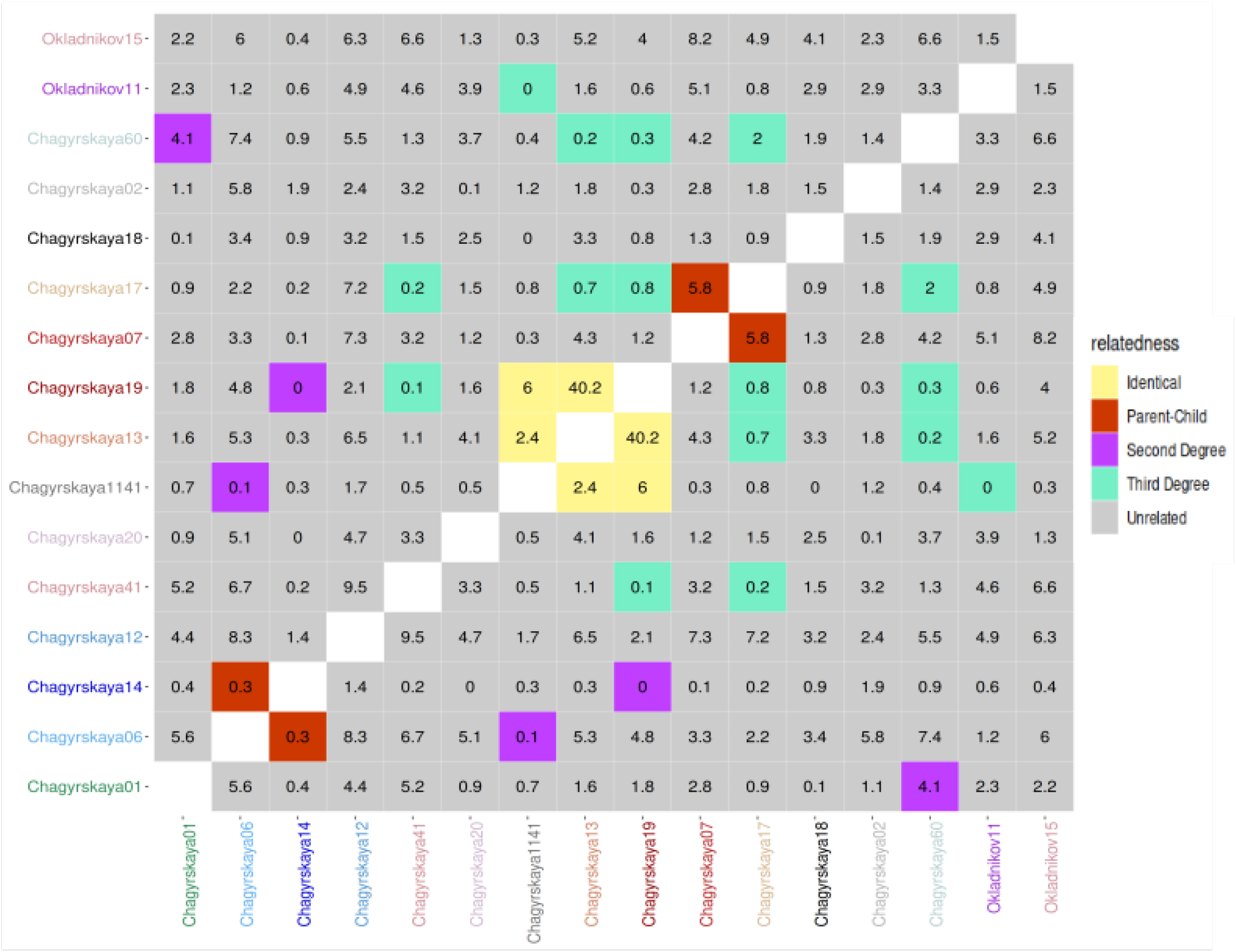
Application of KIN to Neandertal remains from Chagyrskaya and Okladnikov Caves. The color of a square represents the relatedness, while the number within denotes log likelihood ratio (∆*LL*) between the two maximum likelihood models.

Our estimates are consistent with those obtained using READ, except that READ is unable to detect 3rd-degree relationships, and does not distinguish between sibling and parent-child relationships. We also find that both KIN and READ classify Chagyrskaya06 and Chagyrskaya14 as parent-child with low confidence, but from the morphology they likely stem from the same individual. We believe that the low confidence mis-classification may be due to uncorrected non-human contamination present in these libraries (1% and 2.9% respectively [41]), biasing the estimated differences between the individuals to higher values.

We compared the IBD estimates obtained using KIN to those from lcMLkin for all pairs for whom READ results (with s.d. >1) and KIN results (∆*LL* >1) match (Fig. S10). We find that lcMLkin creates four different clusters based on the coefficient of relatedness (*r*) and the proportion of the genome in the unrelated state (*k*_0_) for different relatedness cases (identical, parent-child, 2^*nd*^ degree and unrelated). However, the location of these clusters strongly deviate from the expected values, likely due to the low coverage and ROH. Remains from the same individual, for example, are expected to be at *k*_0_ = 0 and *r* = 1, but are at *k*_0_ ≈ 0.3, *r* ≈ 0.6 for lcMLkin (Fig. S10).

#### 3.2.1 Ancient modern humans

We further applied KIN to a genome-wide dataset of 118 ancient individuals from the Lech Valley [28]. We compared our relatedness estimates to those obtained by the authors using READ and lcMLkin. We found that KIN was able to confidently classify 85% of the 6903 possible comparisons (∆*LL* > 1), READ 94% (*s.d.* > 1) and lcMLkin 53% of pairs ([28]).

Only for 28 pairs, KIN has confident classifications that differ from those obtained using READ (table S2, Fig. S11). Twenty of these comparisons, KIN predicts to be third-degree relationships, and READ concordantly classifies them as unrelated (as it only infers first and second degree relationships). lcMLkin predicts 3^*rd*^ - 5^*th*^ in 14 of these cases, unrelated in 1 case (see Fig. S11), second degree in 1 case (see Fig. S11), and does not have enough data for classification in 4 cases. For 10 additional pairs, lcMLkin predicts 3^*rd*^ – 5^*th*^ degree, but KIN infers them to be unrelated. There are a total of 83 pairs where READ obtains a confident call (*s.d.* > 1), but KIN does not (∆*LL* < 1). In 80 of these cases, READ classifies them as unrelated, KIN classifies 3^*rd*^ degree, while lcMLkin predicts 3^*rd*^ – 5^*th*^ degree. The remaining cases READ classifies unrelated, while KIN and lcMLkin call a 2^*nd*^-degree relationship. Thus, the vast majority of differences can be explained by READ not considering 3rd-degree relationships.

For less than third-degree relations, only eight cases are classified differently between KIN and READ. lcMLkin matches KIN’s prediction in two cases, and does not have enough data in 5 cases. In the last case, all three methods differ (table S3, Fig. S11), and the true relatedness is unresolved in this case. Finally, there are three disagreements between KIN and lcMLkin in classification of parent-child versus siblings (READ predicts first degree for all three pairs). We plotted the pairwise differences for these three pairs in Fig. S12, and found that the proportion of differences along the genome aligns with the prediction of KIN.

## 4 Discussion

Here, we present a new method called KIN to estimate genetic kinship and the location of IBD tracts from low-coverage data, in presence of long ROH, contamination, as well as ascertainment bias. Our method utilizes a set of HMMs to estimate IBD tracts and uses them to classify each pair of individuals into a possible relatedness case, along with a measure of classification confidence (Fig. S1). We evaluated the method performance of KIN, and compared it to that of READ using simulated pedigrees. Finally, we show applications of KIN on two ancient datasets.

For detecting ROH, there is only one method available that works with low coverage ancient DNA [39]. This software can infer ROH at coverages ≥ 0.3*x*, but requires a large reference panel. Instead, our method detects long ROH based on just the expected heterozygosity *p*_0_, which can be estimated from a small number of unrelated individuals. On simulations, we find that our method reliably inferrs ROH regions with lengths on the order of 10cM from samples sequenced to coverage ≥ 0.1*x* . In our simulations, we find that adding long ROH (≈ 17%) to simulations slightly improves the power of both KIN and READ, particularly at lower coverages. This is likely because ROH actually reduces the variance in differences depending on the relatedness, and makes it easier to correctly classify relatedness. For READ, the presence of ROH causes a bias towards inferring closer relatedness cases, but the model we use in KIN reduces this bias (see Fig. 5).

We show that KIN reliably detects IBD tracts for ≥ 0.1*x* coverage even in the presence of contamination and ascertainment bias, and the accuracy for IBD detection reduces with lower coverages. Adding ROH to the simulations adversely affects IBD prediction in siblings (Fig. S9), but this does not affect the power to correctly classify siblings (Fig. 5).

Contamination can affect the accuracy of relatedness inference for two reasons. For one, if a substantial fraction of samples is contaminated, then the estimation of *p*_0_ becomes inaccurate, because the majority of pairwise differences will include at least one contaminated sample. The second issue is that contaminated samples will look less similar to other individuals, and thus cause a bias towards inferring them to be less related to other individuals. The amount of contamination does not need to be large for this to be important, in our simulations we find that even at contamination levels ≤ 3%, the performance of READ is substantially reduced. When contamination rates are correctly inferred, the correction we implemented leads to improved performance compared to naive methods such as READ, although they too could be ameliorated in a similar way [41]. However, in many cases contamination estimates may be uncertain or inaccurate, and we show in fig. S13 that KIN’s performance is robust to small deviations (small compared to average pairwise heterozygosity) in contamination estimates.

The Lech Valley data has low contamination, and no ROH. For pairwise comparisons with large numbers of overlapping sites (> 10000), KIN, READ and lcMLkin all mostly agree. However, KIN is able to differentiate between parent-child and siblings, and identify second degree relationship from just a few thousand polymorphic sites (≈ 4000) overlapping between samples. KIN can also infer third degree relation with ≈ 30,000 overlapping polymorphic sites. We show that when applied to Neandertal specimens from Chagyrskaya and Okladnikov Caves, KIN identifies a pair of 1^*st*^-Degree relatives as parent-child, which is in agreement with the finding that the mtDNA haplotypes differ between the samples (one sample is male and the other female) [41]. In addition, KIN identifies a pair of 3^*rd*^ degree relatives. In this case of a population with large amounts of ROH, we find that the inference by lcMLkin are heavily biased, but KIN’s model takes ROH into account and both the coefficient of relatedness and *k*_0_ are very close to what would be expected from the inference by both READ and KIN.

One limitation of our approach is that it assumes a single population. In case of a highly structured population, KIN may show inaccurate inference of *p*_0_ causing inaccurate relatedness inference. Also, our method makes the assumption, that the median pairwise genetic difference in the population reflects the population diversity *p*_0_, which fails if almost all individuals in the dataset are related. We may get around this problem by using an estimate of *p*_0_, calculated from known a pair of unrelated individuals from same population, or another population with similar diversity. We provide the user with an option to give an estimate of *p*_0_. The current implementation of KIN is restricted to six relatedness cases we expect to be most common, but it might be feasible to extend it to other cases, such as double first cousins, using a corresponding IBD state transition matrix.

While we have focused on the application of KIN on ancient human samples, the model is not tied to this system. Assuming we know the recombination rate, and hence can estimate the transition matrix (see section 5), KIN can be widely applied to any diploid species. In addition, the output of KIN is a table which shows for each pair, the most likely model, and the second best guess, along with a confidence level represented by the log likelihood ratio. This makes KIN easy to automatize for large datasets. To make application of KIN user-friendly, we provide a python package (KINgaroo) to create input files for KIN from processed bam files, while optionally estimating ROH, and correcting for contamination estimates.

## 5 Materials and methods

### 5.1 Simulations

We use simulations both for estimating the transition matrices and for testing and validating our algorithm. All simulations are performed in a scenario mimicking the analysis of a Neandertal population contaminated by modern humans [26]. We simulate unrelated individuals using msprime [17], followed by an additional step where we simulate related individuals using a predetermined pedigree (Fig. S3).

#### Simulating pedigrees

For our simulations of background diversity, we form a population (Pop1) with constant effective size of 10,000 and sample eight diploid individuals (each made up of two haploid individuals) from 2500 generations ago (Fig. S3). For each individual, we simulate 22 chromosomes with length *L* ≈ 96 Mb (same as chromosome 13) and a recombination rate of *r* = 10^*−*8^ per base pair per generation. We introduce mutations using an infinite sites model with rate *μ* = 10^*−*8^ per base pair per generation.

For the pedigree simulations, we first simulate a recombined set of chromosomes for either parent, and combine them to create the progeny. There are two different ways in which we generate recombination points. For the estimation of transition probabilities, we simulate recombination by first drawing the number of breakpoints as a Poisson random variable with parameter *rL*, and use a uniform distribution on [1, *L*] to sample the positions of recombination points. For the testing of our method, we use Ped-sim [6] to simulate recombination points. This allows us to take into account sex-specific recombination rates and crossover interference, and thus is expected to give a more realistic recombination landscape. The pedigree simulations result in nine addition individuals, resulting in a total sample of 17 individuals.

#### Transition matrices

KIN requires a transition matrix for each relatedness case, which we estimate by counting the transition between IBD states for all pairs of individuals with that relatedness in a training set of 1000 simulations from our pedigree (Fig. S3). For two cases, siblings and grandparent-grandchild, it is possible to write down the theoretical expectation of the transition matrix:

For grandparent-grandchild, rate matrix is

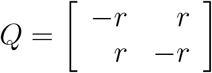

Similarly, for a pair of siblings, we calculate

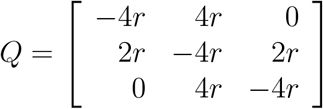

The state space includes all IBD states in this case. For these two cases, we get the transition matrix in each case as *e^Qb^*, where b is the window size.

#### Simulations for method evaluation

Apart from the related and unrelated individuals in Pop1, we simulated more haploid individuals in three other populations to create scenarios with ascertainment bias and contamination (Fig. S3). We simulated two individuals to form an individual each from two other populations (Pop2, Pop3) with split time of 3500 and 4500 generations with Pop1, and sampling time of 2000 generations and 4000 generations ago respectively. We identified the sites that were polymorphic among these two individuals, and used these sites to ascertain the genomes of individuals from Pop1. This scenario roughly models the ascertainment of the Chagyrskaya and Okladnikov Caves data. We tested the performance of our method in presence of long ROH (~ 17%), by simulating regions of homozygosity in unrelated individuals with a Markov chain using the transition matrix:

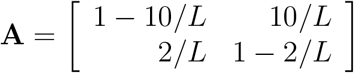

It is worth noting that we introduce long ROH in unrelated individuals, before artificially mating them to form pedigrees, which means that we do not directly introduce ROH in progeny, but it still affects relatedness inference among progeny as shown in Fig. 1. From the steps described above, we got genotypes of individuals in Pop1 in presence/absence of ROH and ascertainment. We further simulated five diploid individuals from Pop4 with split time of 20,000 generations with Pop1, sampled from the present time, as a source of contamination. For cases including contamination, we contaminated eight individuals with varying amounts between 0.5% to 3% contamination, while the remaining nine individuals did not have any contamination. We generated reads (derived/ancestral) for different genomic coverages ranging from 4x to 0.03x, assuming a Poisson distribution.

For testing, we replicate our simulation 60 times, and create data from the same base simulations at varying levels of coverages and different scenarios: We have a control scenario (without ascertainment bias, ROH or contamination), and individual scenarios where we add ROH, SNP ascertainment and contamination. For the evaluation of the ROH-HMM, we also combine ROH with ascertainment or contamination.

## 6 Acknowledgements

We thank Svante Pääbo, Janet Kelso, Harald Ringbauer, Johann Visagie, Zbigniew Jedrzejewski-Szmek, Laurits Skov, Leonardo N. M. Iasi, Alba Bossoms Mesa, Arev P. Sümer, Zuzana Hofmanová, Guido Alberto Gnecchi Ruscone, Ke Wang and Luca Traverso for helpful comments and discussions. This work was supported by the Max Planck Society and the European Research Council (Grant No.: 694707) to Svante Pääbo.

## 7 Contributions

Conceptualization (Design of study): B.M.P.; Software: D.P.; Methodology—lead: D.P.; Methodology—support: B.M.P., S.P.; Formal Analysis: D.P.; Visualization-lead: D.P.; Visualization-support: S.P.; Data Curation: D.P.; Writing—lead: D.P.; Writing—support: B.M.P., S.P.; Supervision: B.M.P.

## 8 Data and material availability

An open-source implementation of KIN and KINgaroo in python along with a toy example dataset, and the scripts to generate our simulations are available on GitHub [9]. We have deposited the version of the software used in the manuscript, along with the above mentioned files on Zenodo [8]. The analysed datasets from Bronze Age Lech Valley were generated in a previous study (ENA, accession no. PRJEB34400). Chagyrskaya and Okladnikov dataset were generated from a study currently in press [41], and the data will be uploaded to European Nucleotide Archive upon publication.

## 9 Ethics declarations

### 9.1 Ethics approval and consent to participate

Not applicable.

### 9.2 Consent for publication

Not applicable.

## 10 Supplementary figures

**Fig. S 7:**
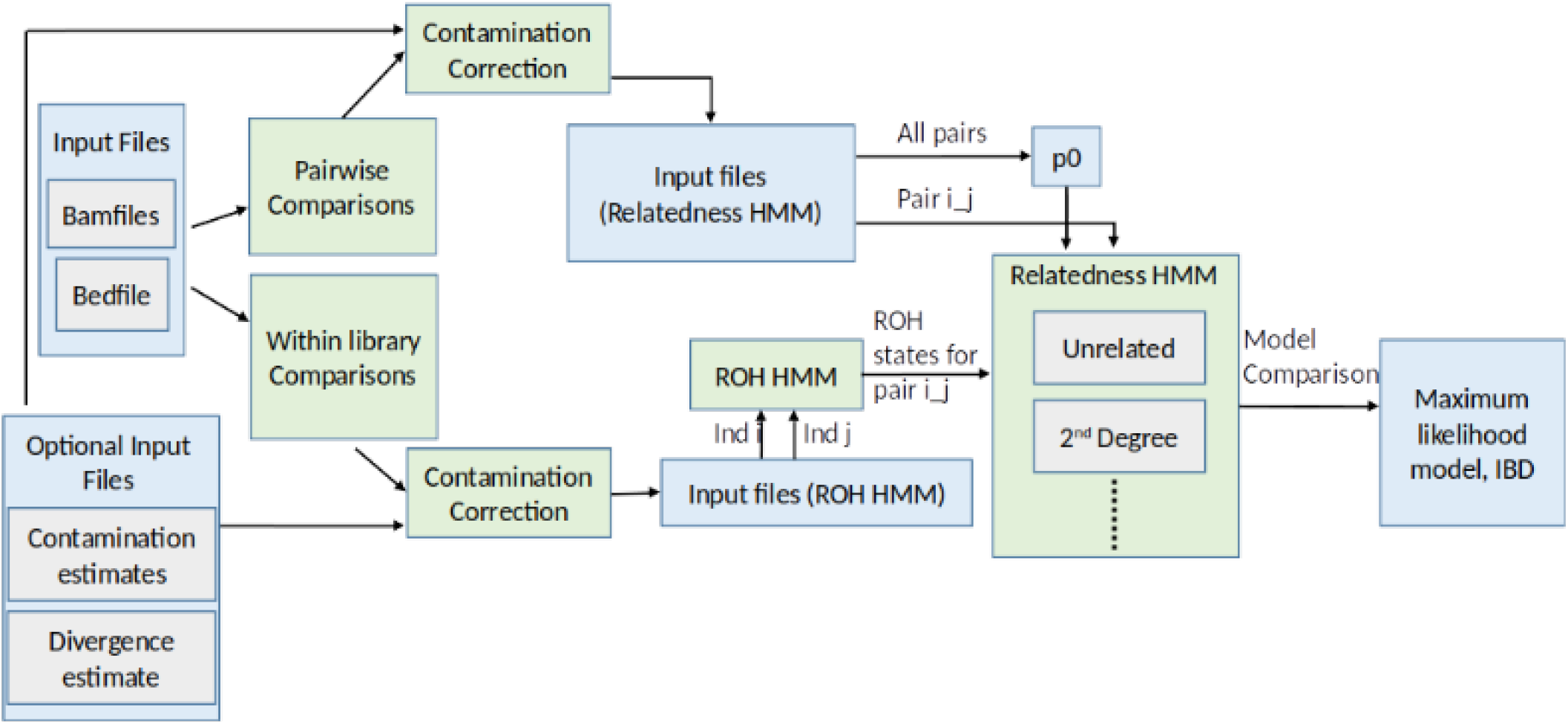
Overall schematic of the method showing the entire pipeline from processed bam files to final relatedness and IBD estimates. Blue boxes show the data files, while green boxes represent scripts.

**Fig. S 8:**
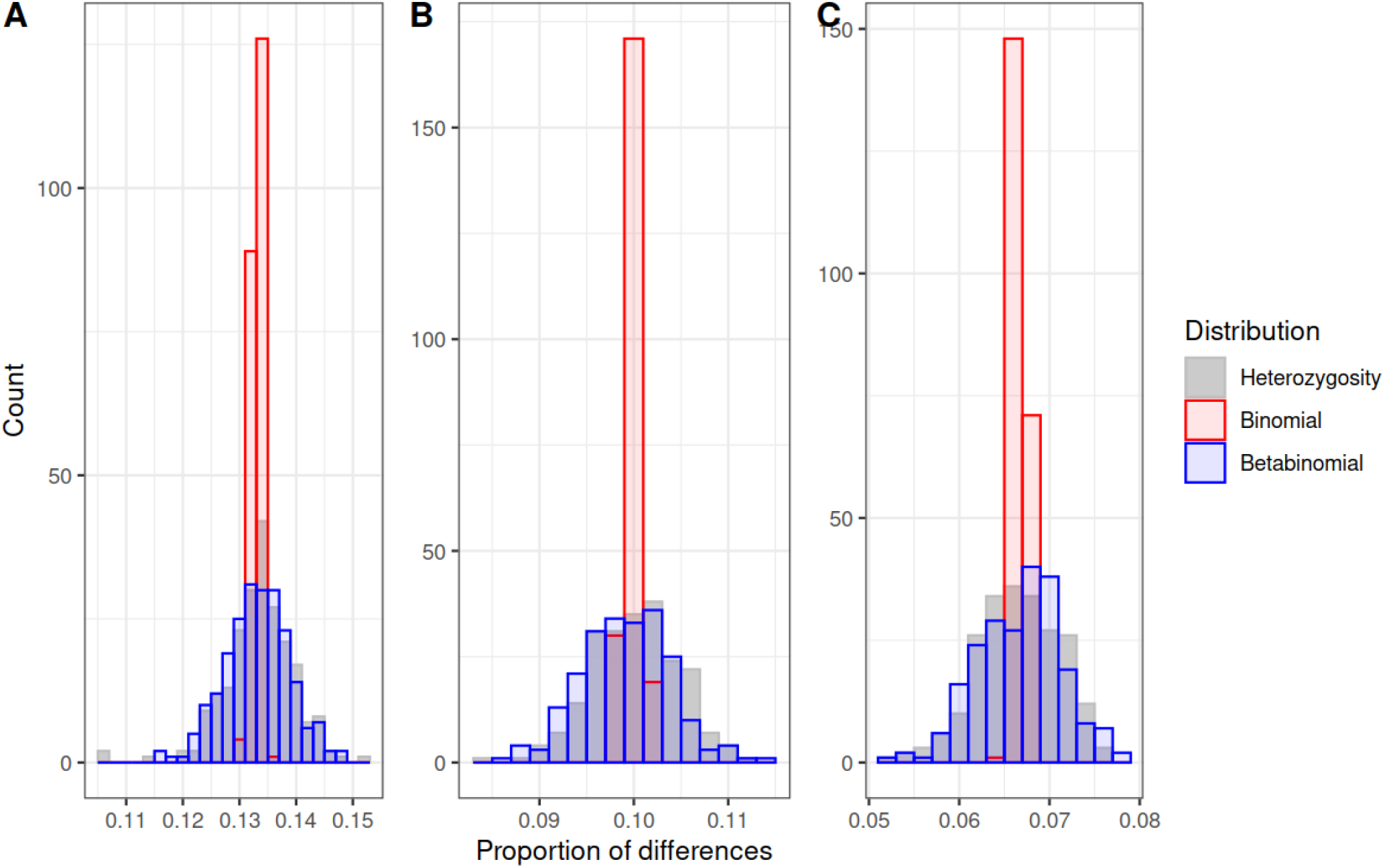
Comparison of fit with Beta-binomial and Binomial distributions. Data is represented by the histogram of proportion of differences from all windows of a (A) unrelated, (B) Parent-Child, (C) Identical pair of individuals. Binomial parameter p is calculated from the data, while Beta-binomial parameters are estimated with corresponding KIN-HMM.

**Fig. S 9:**
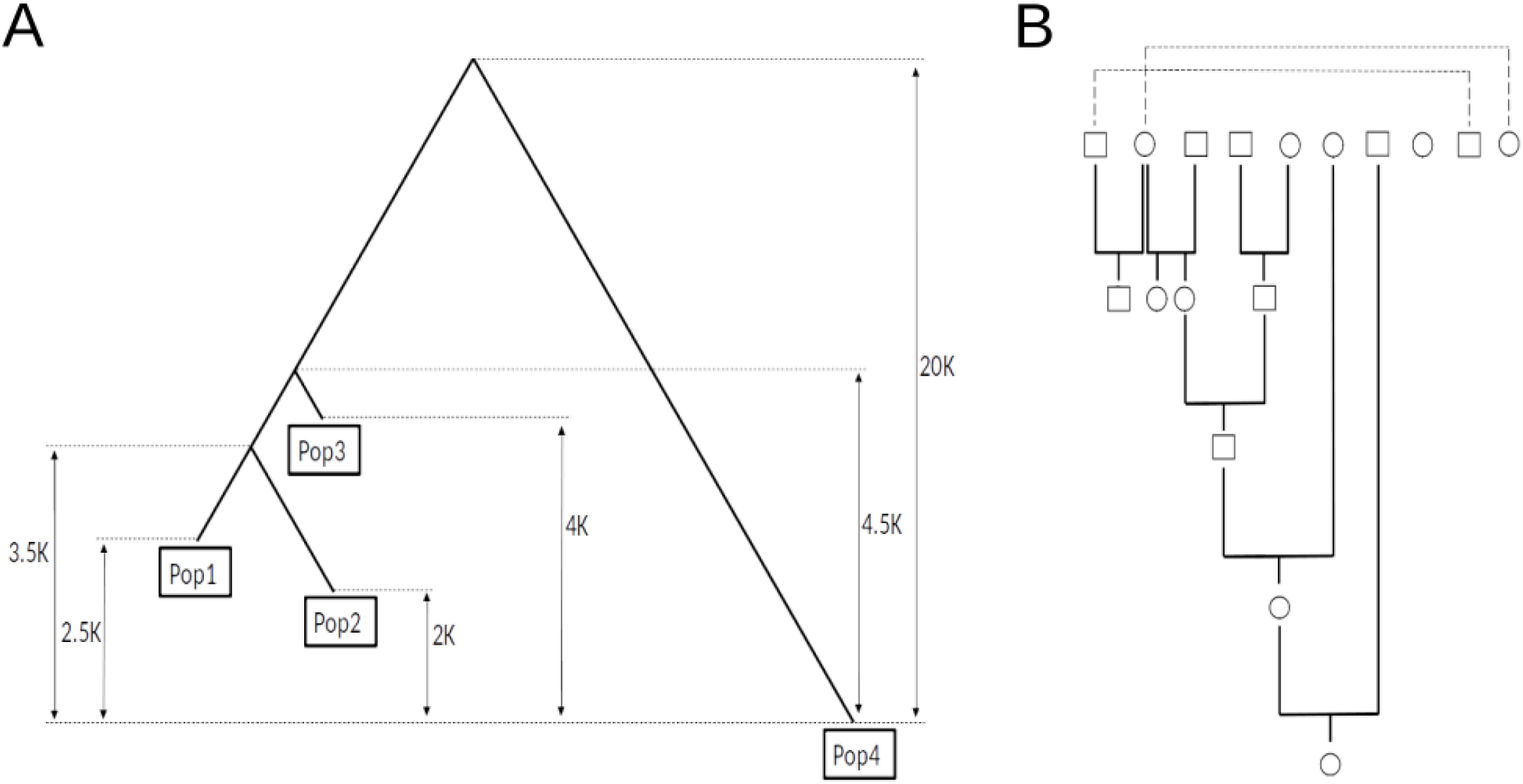
Overview of simulated dataset. (A) We simulated four different populations with split times and sampling times (in generations) mimicking the populations of Chagyrskaya Neandertals, Vindija Neandertals, Altai Neandertals and present day Africans. We artificially mated individuals in Pop1 to create related individuals. We used Pop2 and Pop3 to introduced ascertainment bias in the data, while we simulated contamination from modern humans using Pop4. (B) In Pop1, we sampled 8 diploid unrelated individuals, and artificially mated some of them to make a pedigree with upto 5^*th*^ Degree relatives. Circles represent females, and squares represent males. Dotted lines connect identical pair of individuals.

**Fig. S 10:**
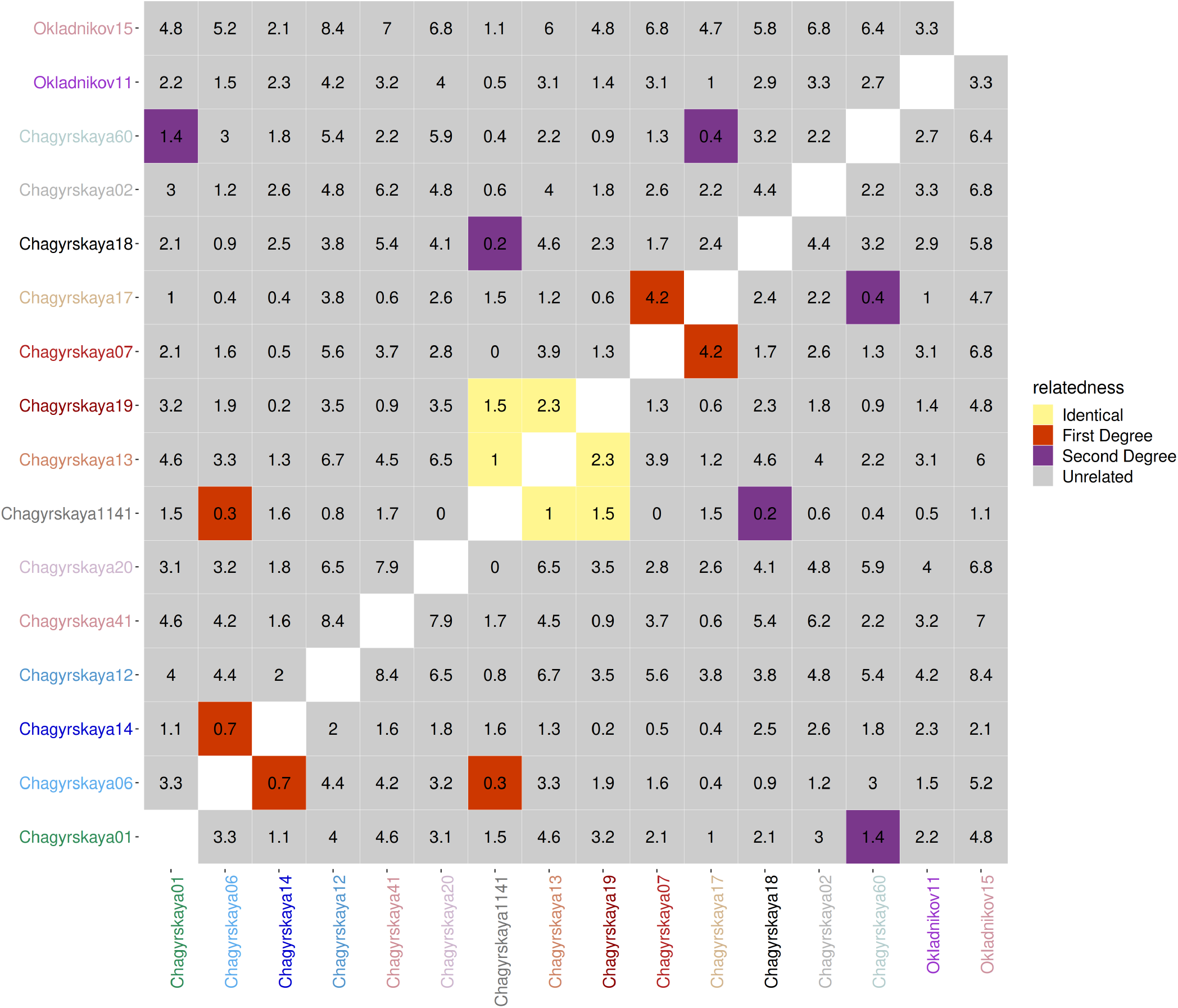
Application of READ on Neandertal specimens from Chagyrskaya and Okladnikov Caves. Color of a square represents the relatedness, while the number denotes standard deviations away from the upper threshold (we show lower threshold for unrelated pairs since upper threshold is not available).

**Fig. S 11:**
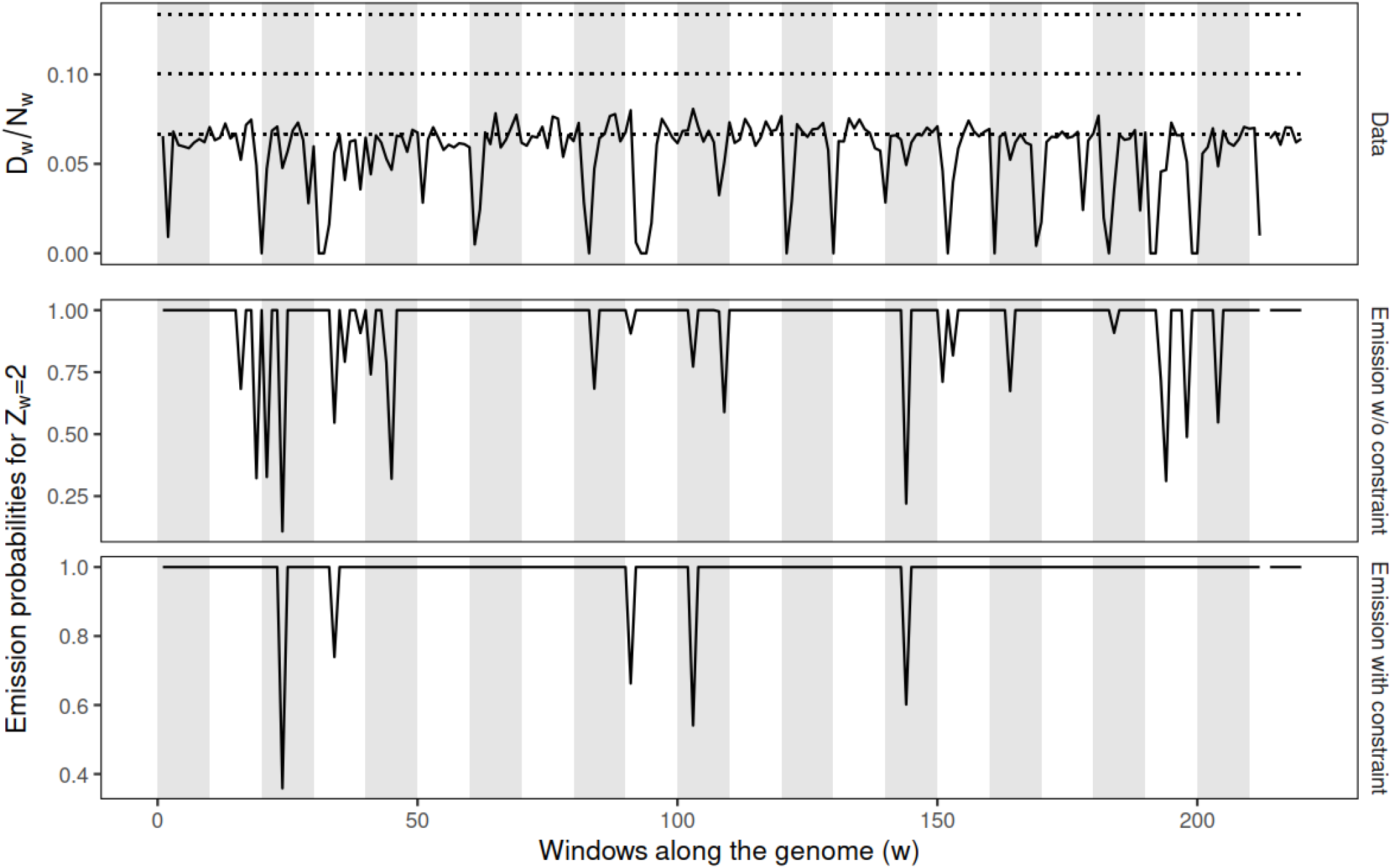
Application of Identical KIN-HMM on a pair of identical individuals with low coverage (0.2x), and ROH tracts. Top panel shows pairwise proportion of differences is close to expectation, but dips in some windows due to presence of ROH tracts. Bottom two panels show *P*(*Data*|*Z_w_* = 2) without constraints and with constraints respectively.

**Fig. S 12:**
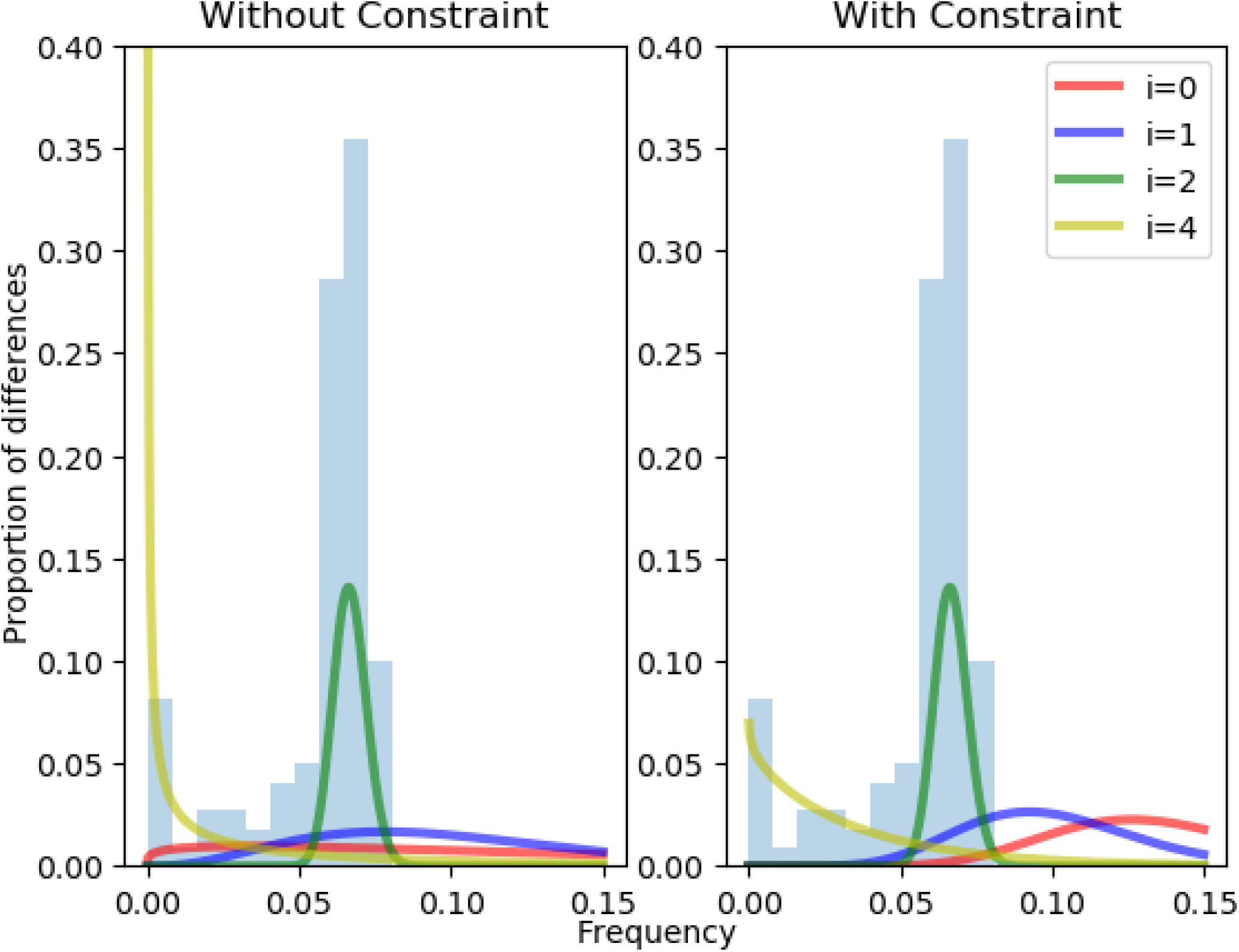
Comparison of beta distributions estimated with the Identical KIN-HMM (left) without and (right) with variance constrained optimization of *δ* parameters.The histogram in each plot shows pairwise differences in genomic windows for identical individuals. Colored lines shows the Beta probability distribution for each *i*, where *i* is the index used for combinations of *Z_w_* and *H_w_*.

**Fig. S 13:**
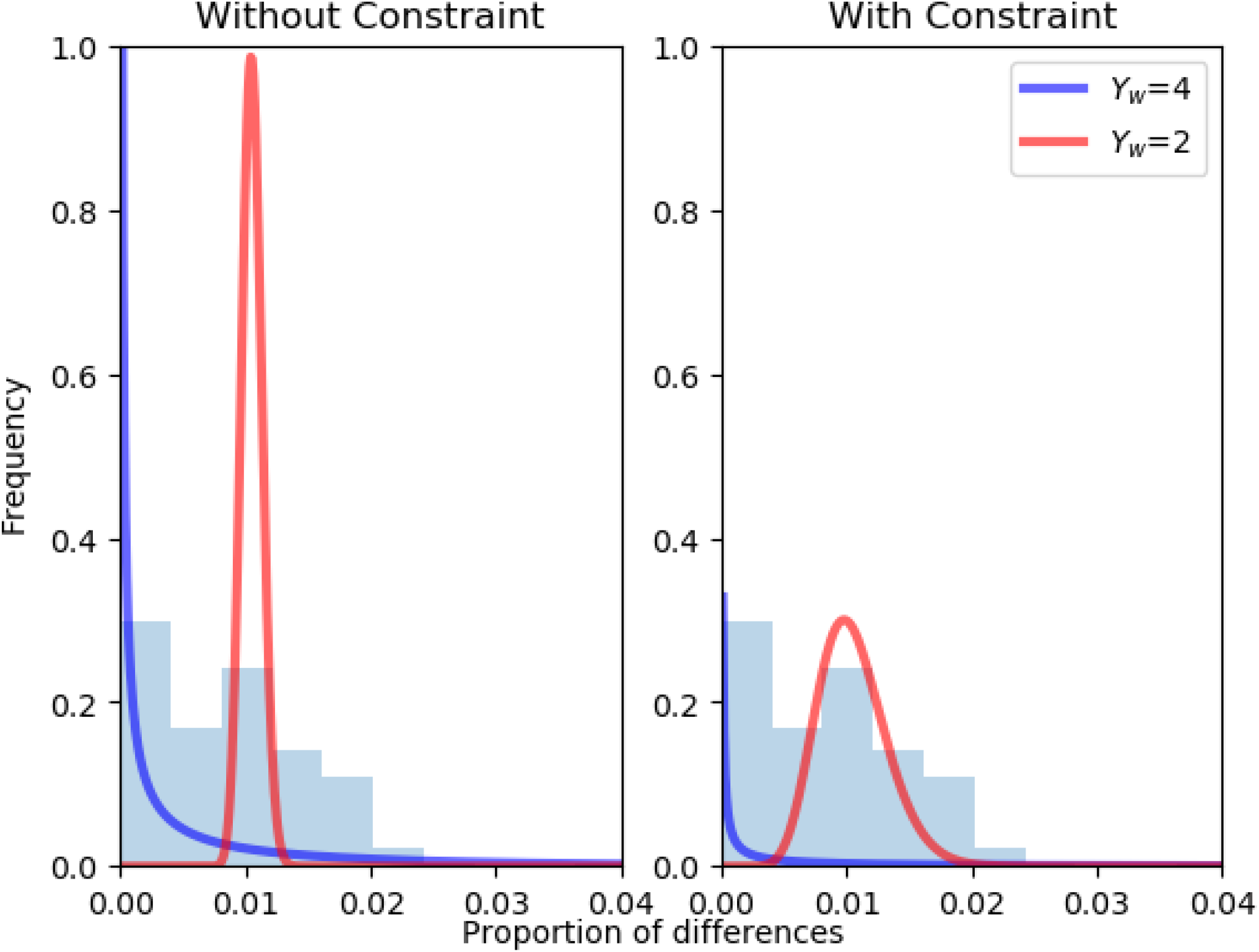
Comparison of Beta distributions estimated with ROH-HMM (left) without and (right) with constrained emissions. The histogram in each plot shows pairwise differences in genomic windows for identical individuals. Colored lines shows the Beta probability distribution for each homozygosity state *Y_w_*.

**Fig. S 14:**
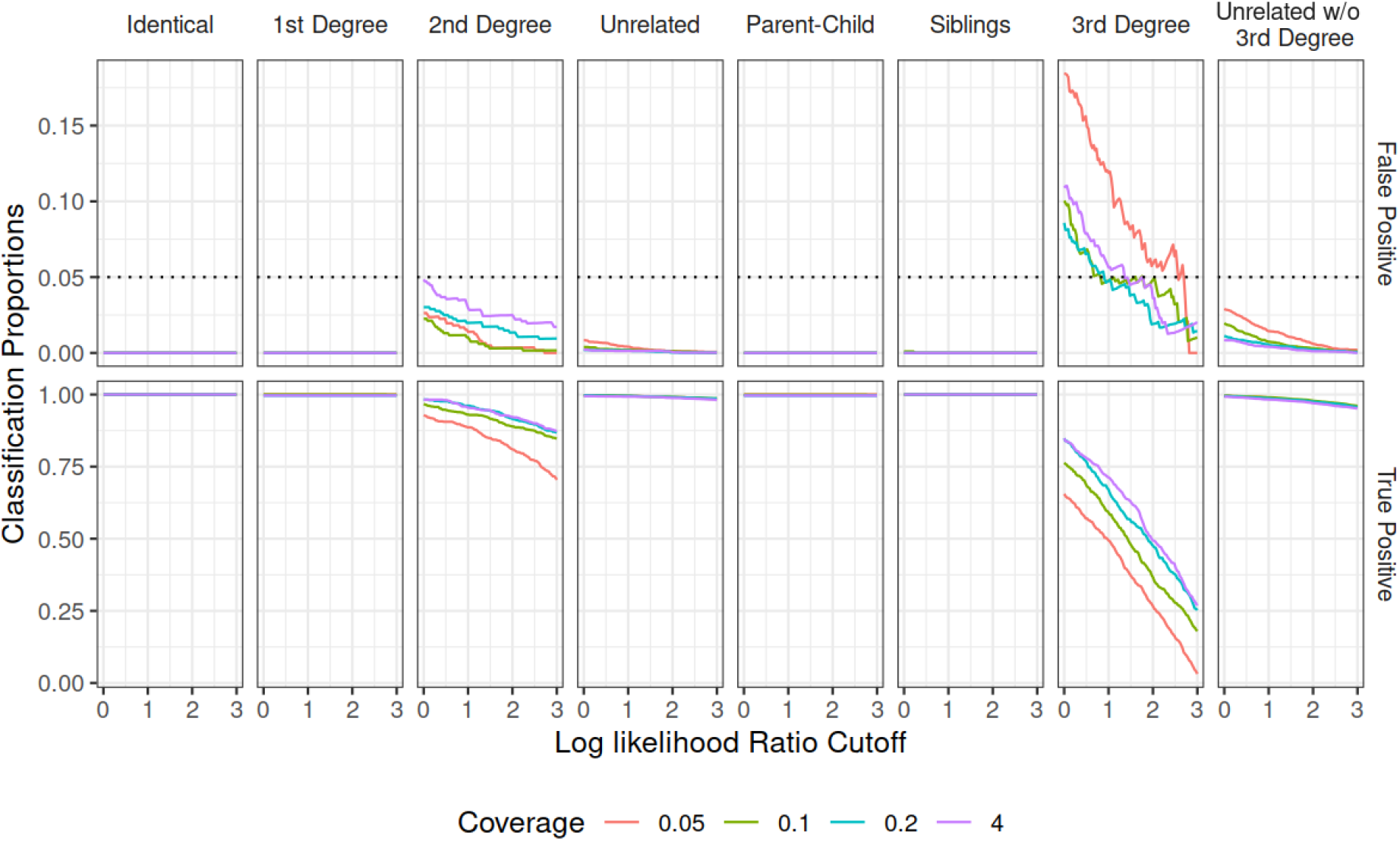
False positive and true positive rates as a function of cutoff on log likelihood ratio. In top panel, dotted line represents 5%. Unrelated label here refers to KIN performance results when all Unrelated, Fifth Degree, Fourth Degree, Third Degree pairs are labelled as Unrelated. ’Unrelated w/o 3^*rd*^ Degree’ refers to the performance results when 3^*rd*^ Degree is classified separately from the unrelated individuals.

**Fig. S 15:**
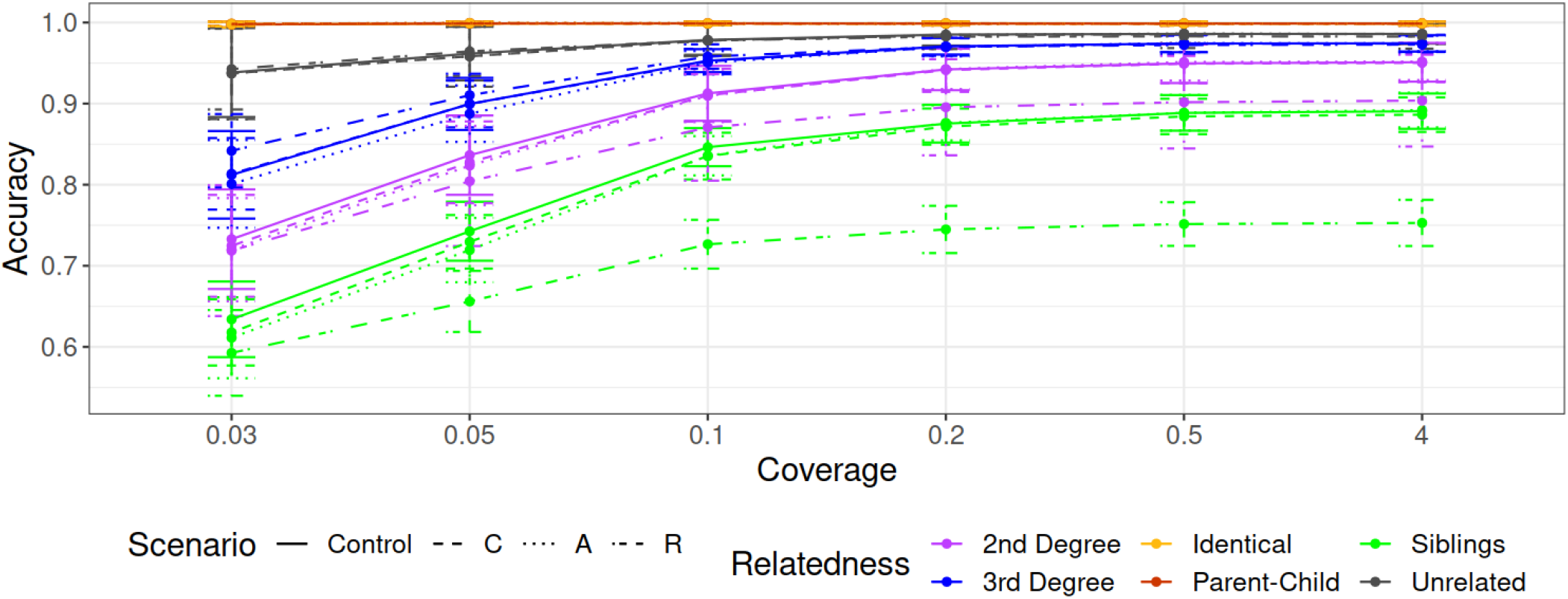
Comparison of IBD estimation in control simulations and in presence of contamination, ascertainment or ROH (denoted in figure legend by Control, C, A and R, respectively) at different coverages. The y-axis shows accuracy calculated over 60 simulations for each relatedness case, for each scenario (control, C, A or R), and six different coverages shown on the x-axis. The error bars display one standard deviation from the mean. Here, accuracy corresponding to relatedness cases for Parent-Child and Identical individuals is always 1, and overlaps with each other.

**Fig. S 16:**
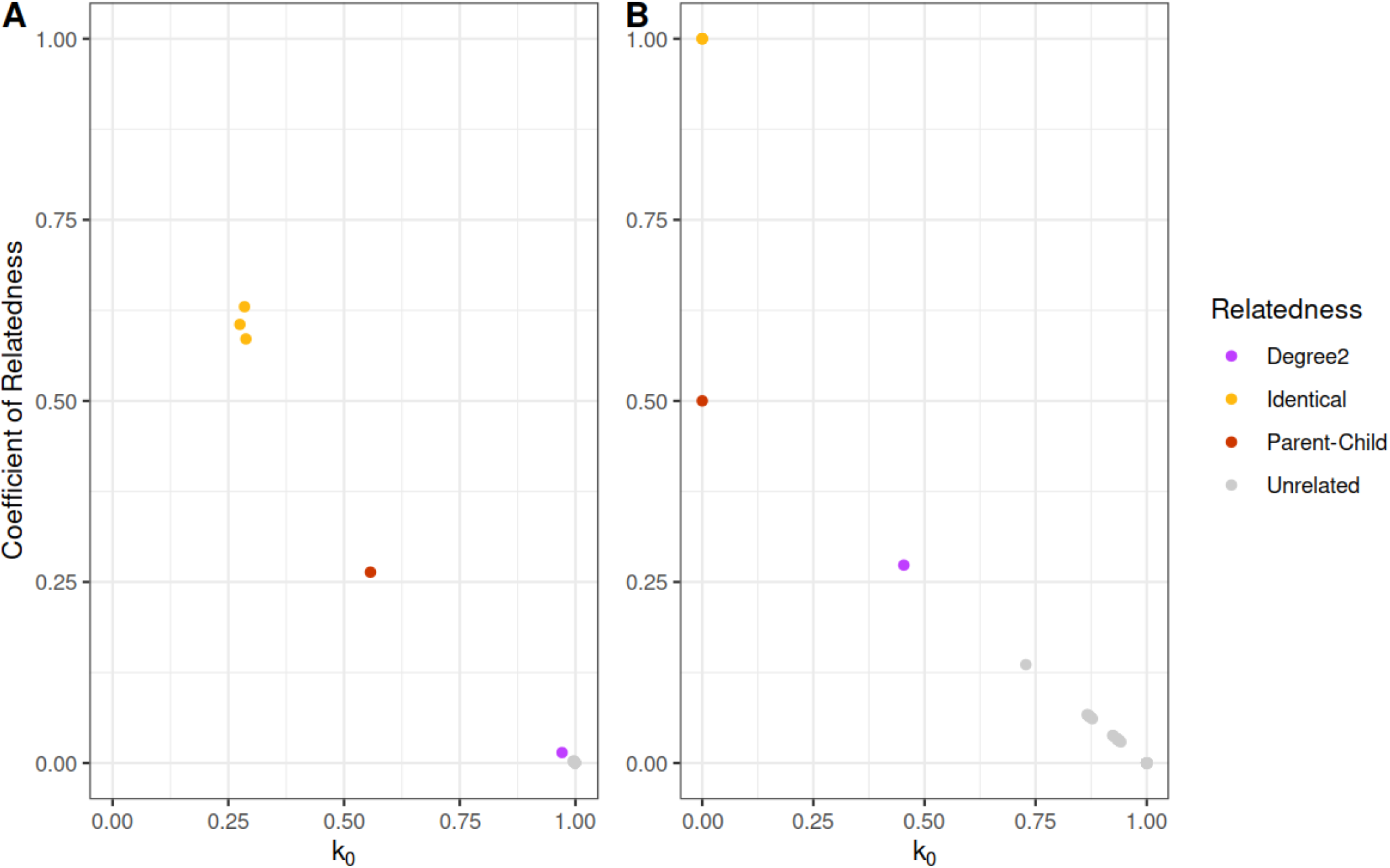
Comparison of IBD states estimated for Chagyrskaya specimens using (A) lcMLkin and (B) KIN. Each plot shows relatedness coefficient (calculated as *k*_2_ + *k*_1_/2) on y-axis plotted against *k*_0_ (proportion of genome with no chromosomes shared). The relatedness shown with different colors is estimated with both READ and KIN.

**Fig. S 17:**
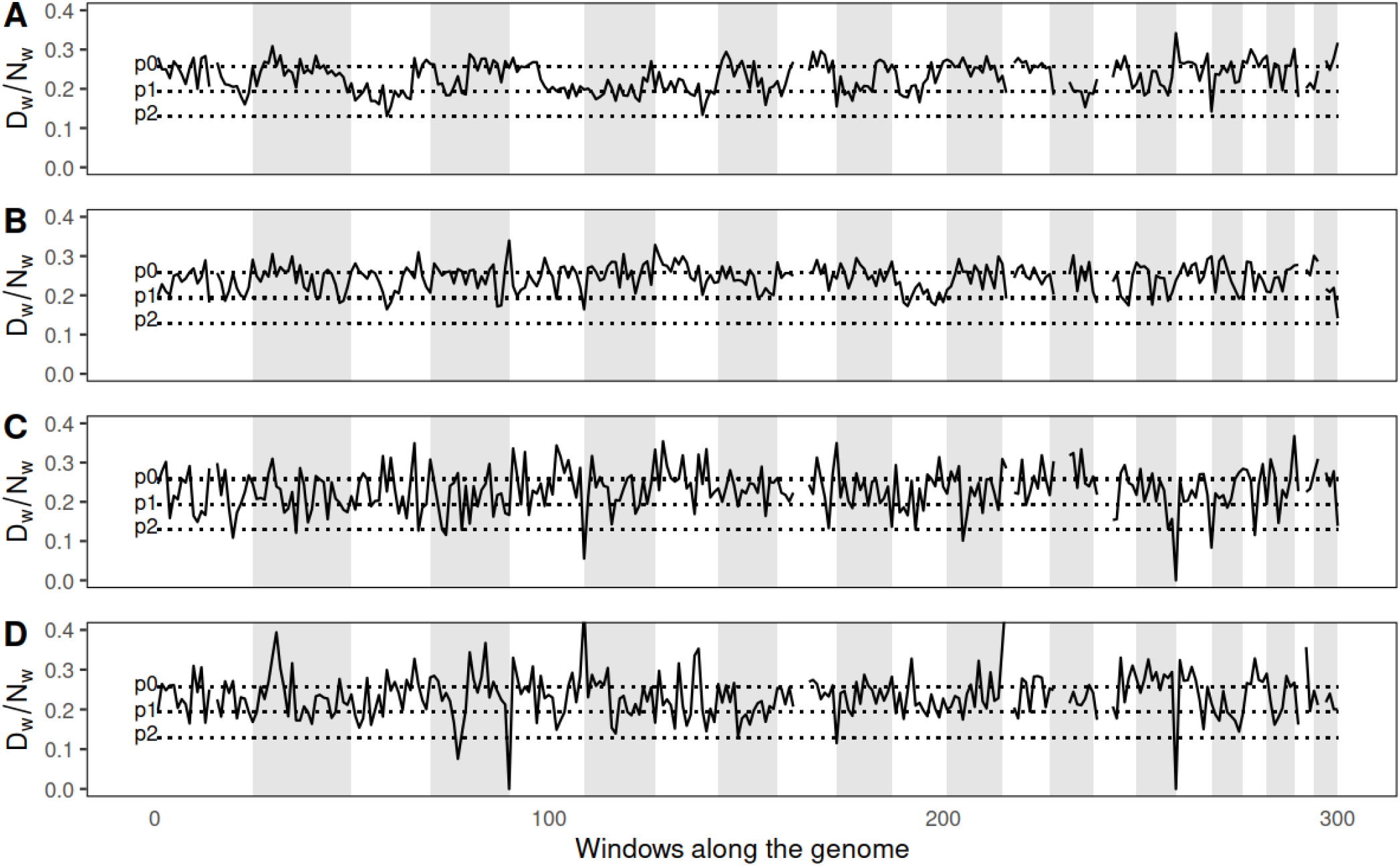
Plots showing proportion of differences in windows along the genome for some pairs of relatives for which there is contradiction among KIN, READ and lcMLkin. (A) shows a case for which KIN and lcMLkin together disagree with READ, and (B) shows a case where KIN and lcMLkin differ, while READ does not classify between contradicting relatedness categories. (C) and (D) show the two unresolved pairs for which all three methods disagree. We expect to see proportion of differences for second degree to be close to *p*_0_ in 50% windows, and *p*_1_ in 50% windows. For third degree we expect 75% windows to be close to *p*_0_, and 25% windows to be close to *p*_1_. For unrelated, we expect the proportion of differences to stay close to *p*_0_ with some noise. (A) AITI95-AITI98 (READ estimates unrelated, lcMLkin and KIN predict second degree. (B) AITI72-AITI77A (READ estimates unrelated, lcMLkin shows unrelated, and KIN predicts third degree). (C) POST131-POST28 (READ estimates Unrelated, lcMLkin shows 3rd-5th degree, and KIN predicts second degree.) (D) ALT3-ALT4 (READ estimates unrelated, lcMLkin shows second degree and KIN predicts third degree.)

**Fig. S 18:**
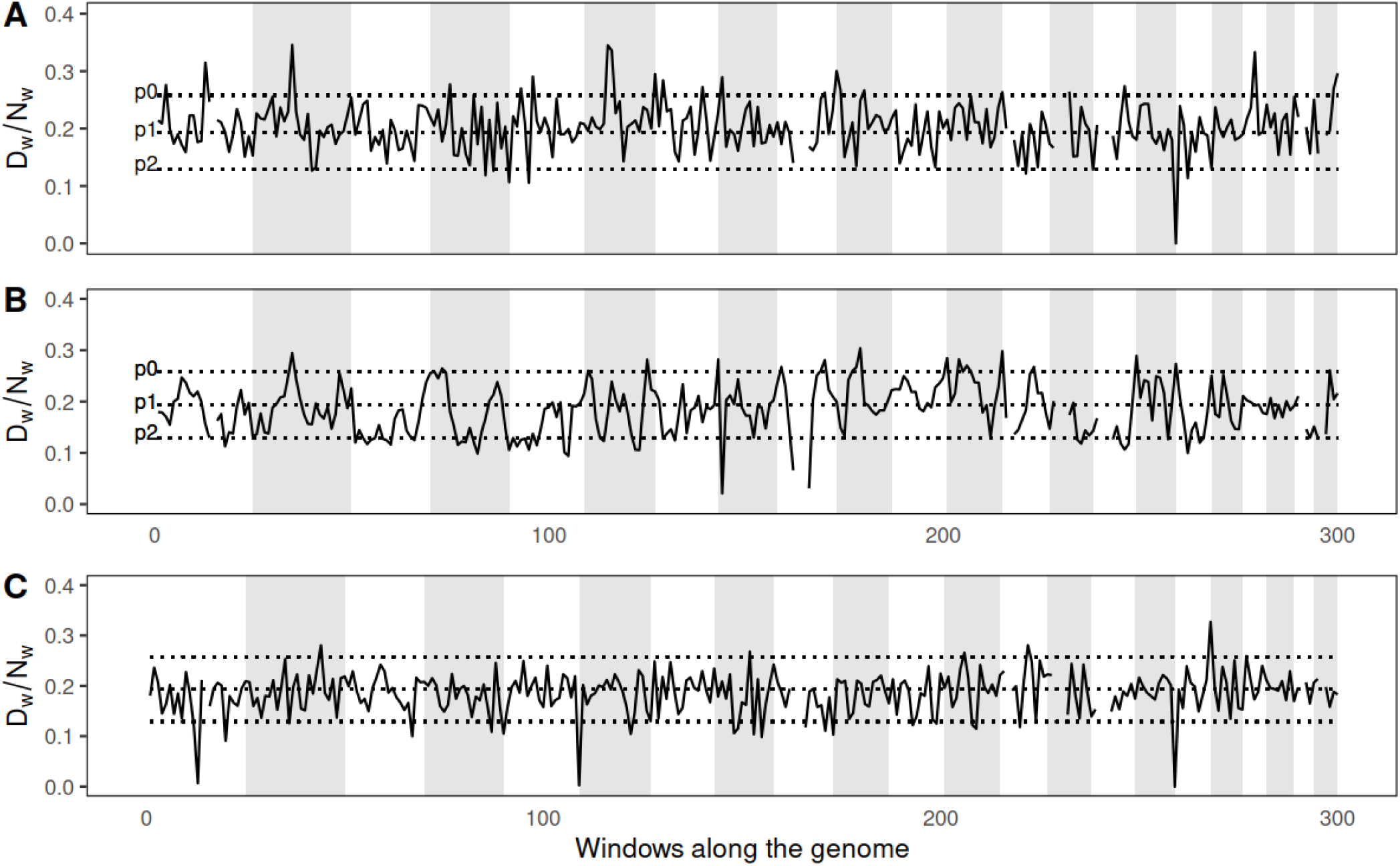
Plots showing proportion of differences in windows along the genome for the first degree relatives for which KIN differs from lcMLkin. We expect to see proportion of differences for siblings to be close to *p*_0_ in 25% windows, *p*_1_ in 50% windows, and *p*_2_ in 25% windows. For parent-child, we expect the proportion of differences to stay close to *p*_1_ with some noise. (A) AITI43-AITI55 (READ estimates first degree, lcMLkin predicts siblings, while KIN predicts parent-child. (B) AITI70-AITI72 (READ estimates first degree, lcMLkin predicts parent-child, while KIN predicts siblings. (C) OBKR76-POST99 (READ estimates first degree, lcMLkin predicts siblings, while KIN predicts parent-child.

**Fig. S 19:**
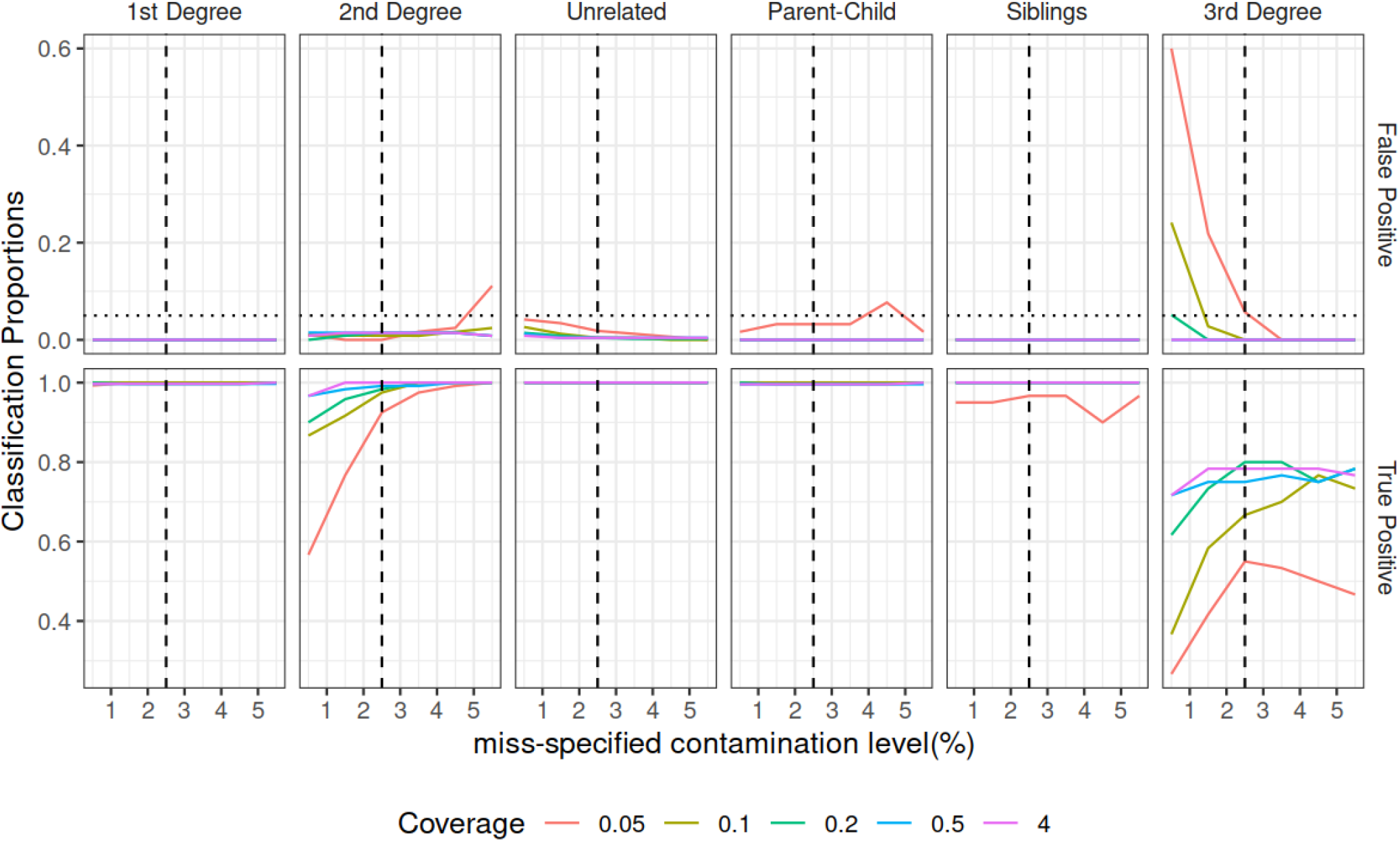
False positive and true positive rates as a function of incorrect contamination estimates provided as input to KIN. Dashed vertical line represents true contamination (2.5%) in simulations, and dotted horizontal line in top panel represents 5% false positive rate. For this analysis we varied the contamination level in one individual from 0.5% to 5.5% while keeping contamination estimates for everyone else constant. We then analyzed the relatedness classification for this individual with all other individuals using 60 runs of simulations (without ascertainment bias or ROH).

## References

[1] Mateusz Baca, Karolina Doan, Maciej Sobczyk, Anna Stankovic, and Piotr Wegleński. Ancient DNA reveals kinship burial patterns of a pre-Columbian Andean community. BMC Genetics, 13(1):30, April 2012.

[2] David J Balding and Richard A Nichols. A method for quantifying differentiation between populations at multi-allelic loci and its implications for investigating identity and paternity. page 10.

[3] Leonard E. Baum, Ted Petrie, George Soules, and Norman Weiss. A Maximization Technique Occurring in the Statistical Analysis of Probabilistic Functions of Markov Chains. The Annals of Mathematical Statistics, 41(1):164–171, February 1970. Publisher: Institute of Mathematical Statistics.

[4] M. Boehnke and N. J. Cox. Accurate inference of relationships in sib-pair linkage studies. American Journal of Human Genetics, 61(2):423–429, August 1997.

[5] Brian L. Browning and Sharon R. Browning. A Fast, Powerful Method for Detecting Identity by Descent. American Journal of Human Genetics, 88(2):173–182, February 2011.

[6] Madison Caballero, Daniel N. Seidman, Ying Qiao, Jens Sannerud, Thomas D. Dyer, Donna M. Lehman, Joanne E. Curran, Ravindranath Duggirala, John Blangero, Shai Carmi, and Amy L. Williams. Crossover interference and sex-specific genetic maps shape identical by descent sharing in close relatives. PLOS Genetics, 15(12):e1007979, December 2019. Publisher: Public Library of Science.

[7] A. P. Dempster, N. M. Laird, and D. B. Rubin. Maximum Likelihood from Incomplete Data via the EM Algorithm. Journal of the Royal Statistical Society. Series B (Methodological), 39(1):1–38, 1977. Publisher: [Royal Statistical Society, Wiley].

[8] Divyaratan Popli. https://doi.org/10.5281/zenodo.7067142.

[9] Divyaratan Popli. https://github.com/DivyaratanPopli/Kinship_inference/releases/tag/v3.1.3, September 2022.

[10] T. Egeland, P. F. Mostad, B. Mevåg, and M. Stenersen. Beyond traditional paternity and identification cases: Selecting the most probable pedigree. Forensic Science International, 110(1):47–59, May 2000.

[11] Richard E. Green, Johannes Krause, Adrian W. Briggs, Tomislav Maricic, Udo Stenzel, Martin Kircher, Nick Patterson, Heng Li, Weiwei Zhai, Markus Hsi-Yang Fritz, Nancy F. Hansen, Eric Y. Durand, Anna-Sapfo Malaspinas, Jeffrey D. Jensen, Tomas Marques-Bonet, Can Alkan, Kay Prüfer, Matthias Meyer, Hernán A. Burbano, Jeffrey M. Good, Rigo Schultz, Ayinuer Aximu-Petri, Anne Butthof, Barbara Höber, Barbara Höffner, Madlen Siegemund, Antje Weihmann, Chad Nusbaum, Eric S. Lander, Carsten Russ, Nathaniel Novod, Jason Affourtit, Michael Egholm, Christine Verna, Pavao Rudan, Dejana Brajkovic, Željko Kucan, Ivan Gušic, Vladimir B. Doronichev, Liubov V. Golovanova, Carles Lalueza-Fox, Marco de la Rasilla, Javier Fortea, Antonio Rosas, Ralf W. Schmitz, Philip L. F. Johnson, Evan E. Eichler, Daniel Falush, Ewan Birney, James C. Mullikin, Montgomery Slatkin, Rasmus Nielsen, Janet Kelso, Michael Lachmann, David Reich, and Svante Pääbo. A Draft Sequence of the Neandertal Genome. Science (New York, N.Y.), 328(5979):710–722, May 2010.

[12] Alexander Gusev, Jennifer K. Lowe, Markus Stoffel, Mark J. Daly, David Altshuler, Jan L. Breslow, Jeffrey M. Friedman, and Itsik Pe’er. Whole population, genome-wide mapping of hidden relatedness. Genome Research, 19(2):318–326, February 2009.

[13] Wolfgang Haak, Iosif Lazaridis, Nick Patterson, Nadin Rohland, Swapan Mallick, Bastien Llamas, Guido Brandt, Susanne Nordenfelt, Eadaoin Harney, Kristin Stewardson, Qiaomei Fu, Alissa Mittnik, Eszter Bánffy, Christos Economou, Michael Francken, Susanne Friederich, Rafael Garrido Pena, Fredrik Hallgren, Valery Khartanovich, Aleksandr Khokhlov, Michael Kunst, Pavel Kuznetsov, Harald Meller, Oleg Mochalov, Vayacheslav Moiseyev, Nicole Nicklisch, Sandra L. Pichler, Roberto Risch, Manuel A. Rojo Guerra, Christina Roth, Anna Szécsényi-Nagy, Joachim Wahl, Matthias Meyer, Johannes Krause, Dorcas Brown, David Anthony, Alan Cooper, Kurt Werner Alt, and David Reich. Massive migration from the steppe was a source for Indo-European languages in Europe. Nature, 522(7555):207–211, June 2015.

[14] D Habier, R L Fernando, and J C M Dekkers. The Impact of Genetic Relationship Information on Genome-Assisted Breeding Values. Genetics, 177(4):2389–2397, December 2007.

[15] Chad D. Huff, David J. Witherspoon, Tatum S. Simonson, Jinchuan Xing, W. Scott Watkins, Yuhua Zhang, Therese M. Tuohy, Deborah W. Neklason, Randall W. Burt, Stephen L. Guthery, Scott R. Woodward, and Lynn B. Jorde. Maximum-likelihood estimation of recent shared ancestry (ERSA). Genome Research, 21(5):768–774, May 2011.

[16] M. Kardos, G. Luikart, and F. W. Allendorf. Measuring individual inbreeding in the age of genomics: marker-based measures are better than pedigrees. Heredity, 115(1):63–72, July 2015. Bandiera abtest: a Cg type: Nature Research Journals Number: 1 Primary atype: Research Publisher: Nature Publishing Group Subject term: Evolution Subject term id: evolution.

[17] Jerome Kelleher, Alison M. Etheridge, and Gilean McVean. Efficient Coalescent Simulation and Genealogical Analysis for Large Sample Sizes. PLOS Computational Biology, 12(5):e1004842, May 2016.

[18] Kseniya A. Kolobova, Richard G. Roberts, Victor P. Chabai, Zenobia Jacobs, Maciej T. Krajcarz, Alena V. Shalagina, Andrey I. Krivoshapkin, Bo Li, Thorsten Uthmeier, Sergey V. Markin, Mike W. Morley, Kieran O’Gorman, Natalia A. Rudaya, Sahra Talamo, Bence Viola, and Anatoly P. Derevianko. Archaeological evidence for two separate dispersals of Neanderthals into southern Siberia. Proceedings of the National Academy of Sciences, 117(6):2879–2885, February 2020. Publisher: National Academy of Sciences Section: Biological Sciences.

[19] Thorfinn Sand Korneliussen, Anders Albrechtsen, and Rasmus Nielsen. ANGSD: Analysis of Next Generation Sequencing Data. BMC Bioinformatics, 15(1):356, November 2014.

[20] Thorfinn Sand Korneliussen and Ida Moltke. NgsRelate: a software tool for estimating pairwise relatedness from next-generation sequencing data. Bioinformatics (Oxford, England), 31(24):4009–4011, December 2015.

[21] Jose Manuel Monroy Kuhn, Mattias Jakobsson, and Torsten Günther. Estimating genetic kin relationships in prehistoric populations. PLOS ONE, 13(4):e0195491, April 2018. Publisher: Public Library of Science.

[22] Hong Li, Gustavo Glusman, Hao Hu, Shankaracharya, Juan Caballero, Robert Hubley, David Witherspoon, Stephen L. Guthery, Denise E. Mauldin, Lynn B. Jorde, Leroy Hood, Jared C. Roach, and Chad D. Huff. Relationship Estimation from Whole-Genome Sequence Data. PLOS Genetics, 10(1):e1004144, January 2014. Publisher: Public Library of Science.

[23] Hong Li, Gustavo Glusman, Chad Huff, Juan Caballero, and Jared C. Roach. Accurate and Robust Prediction of Genetic Relationship from Whole-Genome Sequences. PLOS ONE, 9(2):e85437, February 2014. Publisher: Public Library of Science.

[24] Mikhail Lipatov, Komal Sanjeev, Rob Patro, and Krishna R. Veeramah. Maximum Likelihood Estimation of Biological Relatedness from Low Coverage Sequencing Data. Technical report, July 2015. Company: Cold Spring Harbor Laboratory Distributor: Cold Spring Harbor Laboratory Label: Cold Spring Harbor Laboratory Section: New Results Type: article.

[25] Michael Lynch and Kermit Ritland. Estimation of Pairwise Relatedness With Molecular Markers. Genetics, 152(4):1753–1766, August 1999. Publisher: Genetics Section: Investigations.

[26] Fabrizio Mafessoni, Steffi Grote, Cesare de Filippo, Viviane Slon, Kseniya A. Kolobova, Bence Viola, Sergey V. Markin, Manjusha Chintalapati, Stephane Peyrégne, Laurits Skov, Pontus Skoglund, Andrey I. Krivoshapkin, Anatoly P. Derevianko, Matthias Meyer, Janet Kelso, Benjamin Peter, Kay Prüfer, and Svante Pääbo. A high-coverage Neandertal genome from Chagyrskaya Cave. Proceedings of the National Academy of Sciences, 117(26):15132–15136, June 2020. Publisher: Proceedings of the National Academy of Sciences.

[27] Ani Manichaikul, Josyf C. Mychaleckyj, Stephen S. Rich, Kathy Daly, Michèle Sale, and Wei-Min Chen. Robust relationship inference in genome-wide association studies. Bioinformatics, 26(22):2867–2873, November 2010.

[28] Alissa Mittnik, Ken Massy, Corina Knipper, Fabian Wittenborn, Ronny Friedrich, Saskia Pfrengle, Marta Burri, Nadine Carlichi-Witjes, Heidi Deeg, Anja Furtwängler, Michaela Harbeck, Kristin von Heyking, Catharina Kociumaka, Isil Kucukkalipci, Susanne Lindauer, Stephanie Metz, Anja Staskiewicz, Andreas Thiel, Joachim Wahl, Wolfgang Haak, Ernst Pernicka, Stephan Schiffels, Philipp W. Stockhammer, and Johannes Krause. Kinship-based social inequality in Bronze Age Europe. Science, 366(6466):731–734, November 2019. Publisher: American Association for the Advancement of Science Section: Report.

[29] Erin Murphy. Law and policy oversight of familial searches in recreational genealogy databases. Forensic Science International, 292:e5–e9, November 2018.

[30] Emil Nyerki, Tibor Kalmár, Oszkár Schütz, Rui M. Lima, Endre Neparáczki, Tibor Török, and Zoltán Maróti. An optimized method to infer relatedness up to the 5 ^th^ degree from low coverage ancient human genomes. preprint, Genetics, February 2022.

[31] Pieter A Oliehoek, Jack J Windig, Johan A M van Arendonk, and Piter Bijma. Estimating Relatedness Between Individuals in General Populations With a Focus on Their Use in Conservation Programs. Genetics, 173(1):483–496, May 2006.

[32] Stéphane Peyrégne and Kay Prüfer. Present-Day DNA Contamination in Ancient DNA Datasets. BioEssays, 42(9):2000081, 2020.

[33] Stéphane Peyrégne and Kay Prüfer. Present-Day DNA Contamination in Ancient DNA Datasets. BioEssays, 42(9):2000081, 2020. eprint: https://onlinelibrary.wiley.com/doi/pdf/10.1002/bies.202000081.

[34] Kay Prüfer, Cesare de Filippo, Steffi Grote, Fabrizio Mafessoni, Petra Korlević, Mateja Hajdinjak, Benjamin Vernot, Laurits Skov, Pinghsun Hsieh, Stéphane Peyrégne, David Reher, Charlotte Hopfe, Sarah Nagel, Tomislav Maricic, Qiaomei Fu, Christoph Theunert, Rebekah Rogers, Pontus Skoglund, Manjusha Chintalapati, Michael Dannemann, Bradley J. Nelson, Felix M. Key, Pavao Rudan, Željko Kućan, Ivan Gušić, Liubov V. Golovanova, Vladimir B. Doronichev, Nick Patterson, David Reich, Evan E. Eichler, Montgomery Slatkin, Mikkel H. Schierup, Aida M. Andrés, Janet Kelso, Matthias Meyer, and Svante Pääbo. A high-coverage Neandertal genome from Vindija Cave in Croatia. Science (New York, N.Y.), 358(6363):655–658, November 2017.

[35] Kay Prüfer, Fernando Racimo, Nick Patterson, Flora Jay, Sriram Sankararaman, Susanna Sawyer, Anja Heinze, Gabriel Renaud, Peter H. Sudmant, Cesare de Filippo, Heng Li, Swapan Mallick, Michael Dannemann, Qiaomei Fu, Martin Kircher, Martin Kuhlwilm, Michael Lachmann, Matthias Meyer, Matthias Ongyerth, Michael Siebauer, Christoph Theunert, Arti Tandon, Priya Moorjani, Joseph Pickrell, James C. Mullikin, Samuel H. Vohr, Richard E. Green, Ines Hellmann, Philip L. F. Johnson, Hélène Blanche, Howard Cann, Jacob O. Kitzman, Jay Shendure, Evan E. Eichler, Ed S. Lein, Trygve E. Bakken, Liubov V. Golovanova, Vladimir B. Doronichev, Michael V. Shunkov, Anatoli P. Derevianko, Bence Viola, Montgomery Slatkin, David Reich, Janet Kelso, and Svante Pääbo. The complete genome sequence of a Neanderthal from the Altai Mountains. Nature, 505(7481):43–49, January 2014. Number: 7481 Publisher: Nature Publishing Group.

[36] Kay Prüfer, Udo Stenzel, Michael Hofreiter, Svante Pääbo, Janet Kelso, and Richard E. Green. Computational challenges in the analysis of ancient DNA. Genome Biology, 11(5):R47, May 2010.

[37] Shaun Purcell, Benjamin Neale, Kathe Todd-Brown, Lori Thomas, Manuel A. R. Ferreira, David Bender, Julian Maller, Pamela Sklar, Paul I. W. de Bakker, Mark J. Daly, and Pak C. Sham. PLINK: A Tool Set for Whole-Genome Association and Population-Based Linkage Analyses. American Journal of Human Genetics, 81(3):559–575, September 2007.

[38] Natalie Ram, Christi J. Guerrini, and Amy L. McGuire. Genealogy databases and the future of criminal investigation. Science, 360(6393):1078–1079, June 2018. Publisher: American Association for the Advancement of Science Section: Policy Forum.

[39] Harald Ringbauer, John Novembre, and Matthias Steinrücken. Parental relatedness through time revealed by runs of homozygosity in ancient DNA. Nature Communications, 12(1):5425, September 2021. Number: 1 Publisher: Nature Publishing Group.

[40] Martin Sikora, Andaine Seguin-Orlando, Vitor C. Sousa, Anders Albrechtsen, Thorfinn Korneliussen, Amy Ko, Simon Rasmussen, Isabelle Dupanloup, Philip R. Nigst, Marjolein D. Bosch, Gabriel Renaud, Morten E. Allentoft, Ashot Margaryan, Sergey V. Vasilyev, Elizaveta V. Veselovskaya, Svetlana B. Borutskaya, Thibaut Deviese, Dan Comeskey, Tom Higham, Andrea Manica, Robert Foley, David J. Meltzer, Rasmus Nielsen, Laurent Excoffier, Marta Mirazon Lahr, Ludovic Orlando, and Eske Willerslev. Ancient genomes show social and reproductive behavior of early Upper Paleolithic foragers. Science, 358(6363):659–662, November 2017. Publisher: American Association for the Advancement of Science Section: Report.

[41] Laurits Skov. Genetic insights into the social organization of Neanderthals. accepted.

[42] Christoph Theunert, Fernando Racimo, and Montgomery Slatkin. Joint Estimation of Relatedness Coefficients and Allele Frequencies from Ancient Samples. Genetics, 206(2):1025–1035, June 2017. Publisher: Genetics Section: Investigations.

[43] Timothy Thornton, Hua Tang, Thomas J. Hoffmann, Heather M. Ochs-Balcom, Bette J. Caan, and Neil Risch. Estimating Kinship in Admixed Populations. The American Journal of Human Genetics, 91(1):122–138, July 2012.

[44] Stefania Vai, Carlos Eduardo G. Amorim, Martina Lari, and David Caramelli. Kinship Determination in Archeological Contexts Through DNA Analysis. Frontiers in Ecology and Evolution, 8:83, 2020.

[45] Pauli Virtanen, Ralf Gommers, Travis E. Oliphant, Matt Haberland, Tyler Reddy, David Cournapeau, Evgeni Burovski, Pearu Peterson, Warren Weckesser, Jonathan Bright, Stéfan J. van der Walt, Matthew Brett, Joshua Wilson, K. Jarrod Millman, Nikolay Mayorov, Andrew R. J. Nelson, Eric Jones, Robert Kern, Eric Larson, C. J. Carey, İlhan Polat, Yu Feng, Eric W. Moore, Jake VanderPlas, Denis Laxalde, Josef Perktold, Robert Cimrman, Ian Henriksen, E. A. Quintero, Charles R. Harris, Anne M. Archibald, Antônio H. Ribeiro, Fabian Pedregosa, and Paul van Mulbregt. SciPy 1.0: fundamental algorithms for scientific computing in Python. Nature Methods, 17(3):261–272, March 2020. Number: 3 Publisher: Nature Publishing Group.

[46] Ryan K. Waples, Anders Albrechtsen, and Ida Moltke. Allele frequency-free inference of close familial relationships from genotypes or low-depth sequencing data. Molecular Ecology, 28(1):35–48, 2019. eprint: https://onlinelibrary.wiley.com/doi/pdf/10.1111/mec.14954.

[47] Bruce S. Weir, Amy D. Anderson, and Amanda B. Hepler. Genetic relatedness analysis: modern data and new challenges. Nature Reviews Genetics, 7(10):771–780, October 2006. Number: 10 Publisher: Nature Publishing Group.

